# mmVelo: A deep generative model for estimating cell state-dependent dynamics across multiple modalities

**DOI:** 10.1101/2024.12.11.628059

**Authors:** Satoshi Nomura, Yasuhiro Kojima, Kodai Minoura, Shuto Hayashi, Ko Abe, Haruka Hirose, Teppei Shimamura

**Affiliations:** Division of Systems Biology, Nagoya University Graduate School of Medicine; Laboratory of Computational Life Science, National Cancer Center Research Institute; Japanese Red Cross Aichi Medical Center Nagoya Daiichi Hospital; Department of Computational and Systems Biology, Division of Biological Data Science, Medical Research Laboratory, Institute for Integrated Research, Institute of Science Tokyo

**Keywords:** single-cell RNA sequencing (scRNA-seq), single-cell Assay for Transposase Accessible Chromatin with high-throughput sequencing (scATAC-seq), multiomics, RNA velocity, chromatin velocity, gene regulatory network inference, machine learning, deep learning neural, network deep generative model, variational autoencoder (VAE), differentiation

## Abstract

Single-cell multiomics provides unique insight into the regulatory relationships across different biological layers such as the transcriptome and regulome. However, single-cell multiomics is limited by its ability to capture only static snapshots at the time of observation, restricting the reflection of dynamic state changes orchestrated across modalities. RNA velocity analysis of single cells allows for the prediction of temporal changes in the transcriptome; however, the inferred dynamics cannot be applied across all biological layers, specifically in the regulome. Therefore, to address this limitation, we developed multimodal velocity of single cells (mmVelo), a deep generative model designed to estimate cell state-dependent dynamics across multiple modalities. mmVelo estimates cell state dynamics based on spliced and unspliced mRNA expression, and uses multimodal representation learning to project these dynamics onto chromatin accessibility, inferring chromatin velocity at a single-peak resolution. We applied mmVelo to single-cell multiomics data from a developing mouse brain and validated the accuracy of the estimated chromatin accessibility dynamics. Furthermore, using the estimated dynamics, we identified the transcription factors that are crucial for chromatin accessibility regulation in mouse skin. Finally, using multiomics data as a bridge, we demonstrated that during human brain development, the dynamics of missing modalities can be inferred from single-modal data via cross-modal generation. Overall, mmVelo enhances our understanding of the dynamic interactions between modalities, offering insights into the regulatory relationships across molecular layers.

## 1 Introduction

Single-cell multiomics technologies have emerged as powerful tools for investigating regulatory relationships within individual cells (Vandereyken *et al*., 2023, Baysoy et al., 2023). These technologies facilitate simultaneous measurement of multiple modalities from a single cell, providing a comprehensive view of the complex interactions between different molecular layers, including the transcriptome, proteome and regulome (Hu *et al*., 2016, Angermueller et al., 2016, Hou *et al*., 2016, Cao et al., 2018, Clark et al., 2018, Liu et al., 2019, Rooijers et al., 2019, Mimitou et al., 2019, Xing *et al*., 2020, Plongthongkum et al., 2021, Zhu et al., 2021, Sun et al., 2021, Yan et al., 2021, Swanson et al., 2021, Mimitou et al., 2021, Pan et al., 2022, A. F. Chen et al., 2022, Luo et al., 2022, Rang et al., 2022, Wang *et al*., 2021, Xu et al., 2022). Specifically, methods such as SNARE-seq (S. Chen, Lake), Paired-seq (Zhu *et al*., 2019), SHARE-seq (S. Ma *et al*., 2020), and 10x Genomics Multiome enable the concurrent measurement of gene expression (scRNA-seq; single-cell RNA sequencing) and chromatin accessibility (scATAC-seq; single-cell Assay for Transposase Accessible Chromatin with high-throughput sequencing), thereby advancing our understanding of transcriptional regulatory mechanisms. Importantly, regulatory relationships are inherently dynamic and require the incorporation of temporal information from each modality for accurate estimation. Regulatory networks within cells have been estimated using time-series or ordinary differentiation equations (Gardner *et al*., 2003, Di Bernardo, Gardner, and Collins, 2004, Bansal, Gatta, and di Bernardo, 2006, Wu et al., 2014, Ocone et al., 2015, Matsumoto et al., 2020, and B. Ma, Fang, and Jiao, 2020). However, owing to the destructive nature of sequencing experiments, single-cell data represent static snapshots at the time of measurement, hindering evaluation of the coordinated dynamic changes across modalities (Weinreb *et al*., 2018).

Computational approaches such as trajectory inference and RNA velocity have been developed to capture temporal dynamics in single-cell measurements. Trajectory inference algorithms construct graphs based on cell similarity and assign a pseudotime to each cell, placing the cells along a one-dimensional timeline that represents the progression of a biological process (Trapnell *et al*., 2014, Haghverdi *et al*., 2016, Setty *et al*., 2016, Setty *et al*., 2019, Cannoodt, Saelens, and Saeys, 2016, and Saelens *et al*., 2019). However, for graph construction, trajectory directionality needs to be known and changes in cell states that occur in directions orthogonal to a one-dimensional timeline are not captured. In contrast, RNA velocity estimates the time derivative of the gene expression by linking the abundance of unspliced and spliced mRNA using a mathematical model of splicing kinetics, thereby enabling the prediction of future cell states without prior knowledge and is not confined to a one-dimensional timeline (La Manno *et al*., 2018, Bergen *et al*., 2020). However, predicting dynamics based on RNA velocity models is limited to specific modalities in which a mathematical model can be assumed, and corresponding observations such as unspliced and spliced mRNA can be measured (Bergen *et al*., 2021). Extending these temporal predictions to arbitrary modalities other than RNA, such as chromatin accessibility, remains a significant challenge.

Several studies have considered dynamic changes in chromatin accessibility: by incorporating epigenomic data into a differential equation model of gene expression changes over time, MultiVelo extends the RNA velocity framework (C. Li *et al*., 2023). By adding a differential equation model for changes in chromatin accessibility, the model estimates the dynamics of chromatin accessibility for each gene. However, this aggregates the peak accessibility for each gene, making it difficult to determine the dynamics at the resolution of the single-peak level. Chromatin Velocity uses data from simultaneous measurements of heterochromatin and euchromatin, treating them as the relationship between unspliced and spliced mRNA in the RNA velocity framework, to estimate the dynamics of chromatin accessibility (Tedesco *et al*., 2022). This method can be used to estimate changes in chromatin accessibility at the single-peak level. However, it requires specialized measurement techniques, which limits its general applicability. Additionally, gene expression is not measured simultaneously, limiting our understanding of the regulatory mechanisms across multiple molecular layers. Thus, a method that can simultaneously estimate dynamic changes in gene expression and chromatin accessibility at a single-peak resolution has not yet been developed. This poses a significant challenge for elucidating the detailed mechanisms of gene expression regulation.

To address these challenges, we propose multimodal velocity of single cells (mmVelo), a computational framework designed to estimate cell state-dependent dynamics across multiple modalities using single-cell multiomics data. By utilizing splicing kinetics and multimodal representation learning, mmVelo infers cell state dynamics on joint representations and estimates temporal changes in specific modalities by mapping these dynamics, thereby estimating the temporal changes in chromatin accessibility (chromatin velocity). Focusing on gene expression and chromatin accessibility, we demonstrated that mmVelo accurately estimates chromatin velocity at the single-peak resolution. Additionally, by utilizing the estimated chromatin velocity, we showed that it is possible to predict temporal changes in motif enrichment and identify candidate transcription factors (TFs) that regulate chromatin accessibility. Finally, using multiomics data as a bridge, we illustrated that mmVelo enables integrated analysis of single-modality data and allows for the estimation of dynamics in missing modalities, such as predicting chromatin velocity from gene expression data. mmVelo reveals the dynamic interactions between modalities, providing insights into regulatory relationships.

## 2 Results

### 2.1 mmVelo model

mmVelo uses single-cell multiomics data as input data and outputs the dynamics of each modality (Fig. 1). Let ***a****_n_*, ***u****_n_*, and ***s****_n_* represent chromatin accessibility, unspliced mRNA expression, and spliced mRNA expression in the cell *n*, respectively. Using the encoders for each modality, mmVelo infers the distribution *q*(***z****_n_* | ***a****_n_*, ***u****_n_*, ***s****_n_*), representing the cell state. This distribution is expressed as a mixture of experts from the multivariate Gaussian distributions *q*(***z****_n_* |***a****_n_*) and *q*(***z****_n_* | ***u****_n_*, ***s****_n_*) that are inferred from each modality. Samples from the cell state distribution *q*(***z****_n_* | ***a****_n_*, ***u****_n_*, ***s****_n_*) are then used as input data for a decoder for each modality, outputting the probability distributions *p*(***a****_n_* | ***z****_n_*), *p*(***u****_n_* | ***z****_n_*), and *p*(***s****_n_* | ***z****_n_*) corresponding to each modality. Next, mmVelo infers the distribution *q*(***d****_n_* ***z****_n_*) representing the transition vector ***d****_n_* of the cell state over a small-time interval, using samples from the cell state distribution *q*(***z****_n_* | ***a****_n_*, ***u****_n_*, ***s****_n_*). The transition of the cell state over a short time interval was estimated to be consistent with the RNA velocity *d****s****_n_/dt*, which represents the changes in gene expression described based on splicing kinetics. Finally, mmVelo estimates the temporal changes in each modality by mapping the current cell states (***z****_n_*) and the cell states after the small-time interval (***z****_n_* + *ρ****d****_n_*) using the decoder of each modality.

**Figure 1:**
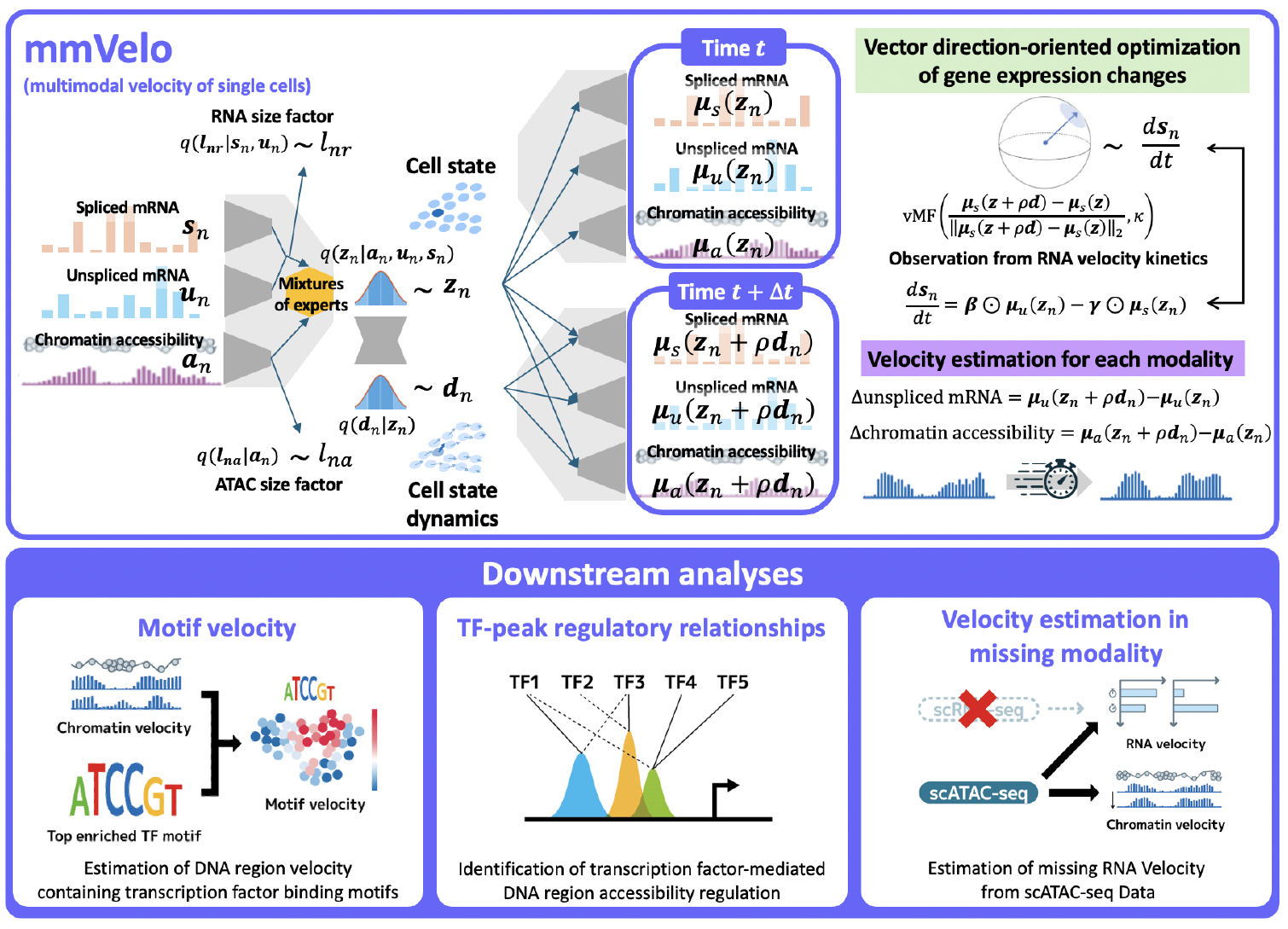
Overview of mmVelo. mmVelo uses single-cell multiomics data as input data and outputs the velocity for each modality. It estimates cell state-dependent dynamics within the cell state space and estimates velocity by mapping these dynamics to the space of each modality. The estimation of cell state-dependent dynamics is conducted based on the assumption of splicing kinetics. Downstream analyses include estimating changes in TF motif accessibility rates, identifying transcription factors that control chromatin accessibility at specific loci, and estimating velocity in a missing modality using single-modality measurement data as input data.

Single-cell multiomics data is difficult to analyze owing to its inherent sparsity and noise compared with single-modality data. This complexity can impede accurate estimation of the differentiation directions in traditional RNA velocity models (S. Ma *et al*., 2020, C. Li et al., 2023). To address this issue, mmVelo integrates multimodal profiles to define cell states, allowing for a more robust estimation of cell states and their neighboring relationships (Minoura *et al*., 2021, Gayoso *et al*., 2015). Furthermore, by modeling RNA velocity along cell state transitions within a low-dimensional manifold, mmVelo accounts for gene-gene coherence, which is not considered in traditional methods that independently solve differential equations for each gene (Nagaharu *et al*., 2022, Lederer et al., 2024, and Gorin *et al*., 2022). To capture the overall direction of gene expression changes, mmVelo employs a von Mises-Fisher distribution for the probability distribution of RNA velocity.

A unique feature of mmVelo is its ability to model chromatin accessibility at the peak level, enabling the estimation of chromatin velocity at a single-peak resolution. This capability allows for dynamic analysis and inference of regulatory relationships at this level. Additionally, by leveraging the temporal changes at the single-peak resolution, mmVelo estimates the dynamics of transcription factor binding motifs at the single-cell level. Another distinctive aspect of mmVelo is its use of multimodal representation learning, which facilitates the prediction of velocities in missing modalities using single-modality data, such as inferring chromatin velocity from scRNA-seq data. Detailed model descriptions and downstream analyses are presented in the Methods section.

### 2.2 mmVelo accurately estimates chromatin accessibility dynamics in embryonic mouse brain

We first applied mmVelo to 10x Multiome data obtained from the embryonic mouse brain on embryonic day 18 (E18). Based on the inferred cell state dynamics and chromatin velocity, the velocity vectors accurately captured the known developmental trajectory of the mammalian cortex (Fig. 2a,b). Specifically, the radial glia cells in the outer subventricular zone are the progenitors of neurons, astrocytes and oligodendrocytes (Merkle *et al*., 2004, Hansen *et al*., 2010, and Pollen *et al*., 2015). Radial glia cells divide into intermediate progenitor cells that give rise to mature excitatory neurons across various cortical layers (Mayer *et al*., 2019). The formation of cortical layers occurs through a process in which neurons migrate in an inside-out manner, with newer cells ascending to the upper layers and older cells remaining in the deeper strata(Nadarajah and Parnavelas, 2002).

**Figure 2:**
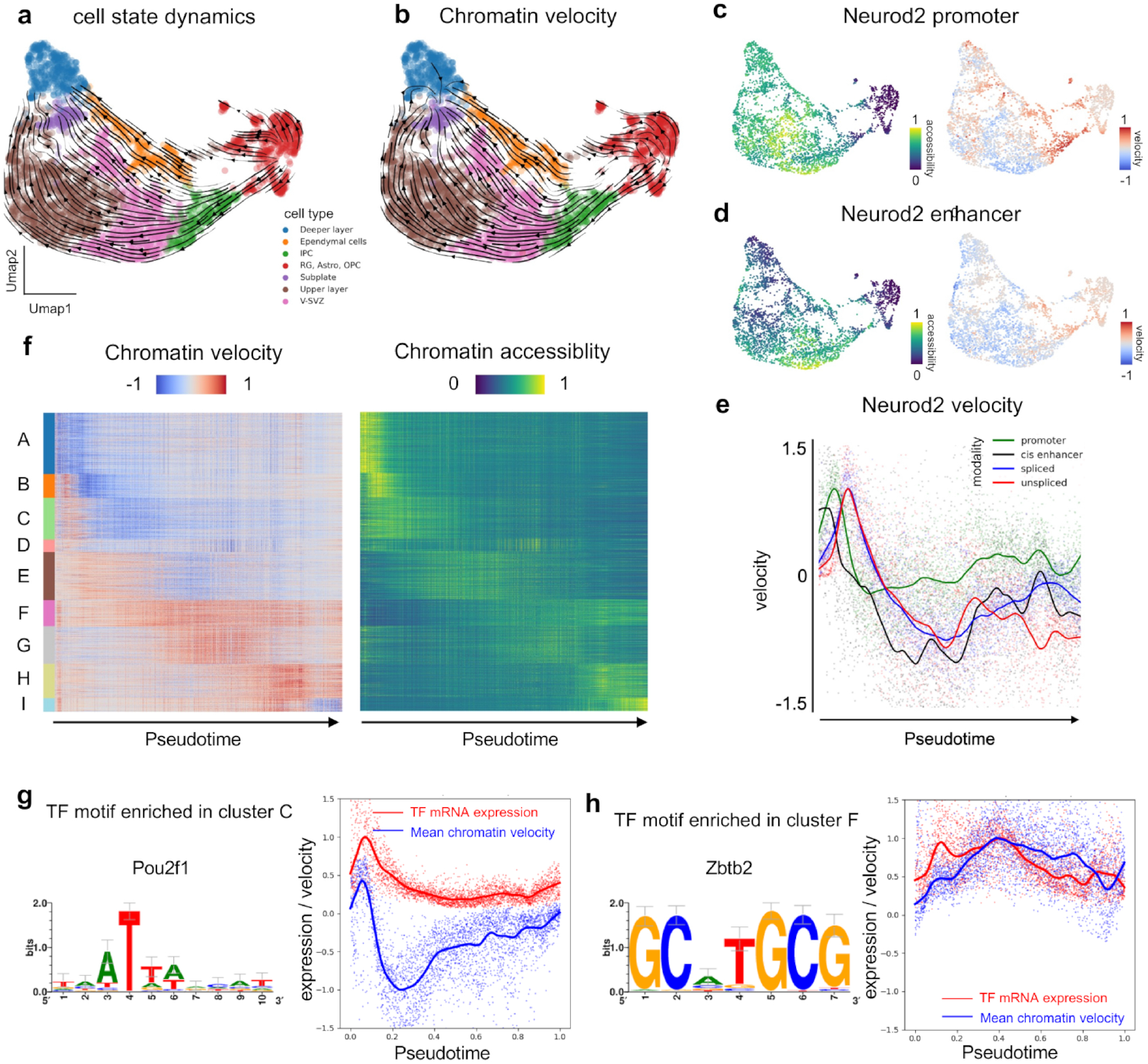
mmVelo accurately estimates chromatin accessibility dynamics in embryonic mouse brain. **a-b**, UMAP coordinates with stream plots of cell state dynamics (**a**) and chromatin velocity (**b**) from mmVelo. IPC, intermediate progenitor cells; RG, radial glia; Astro, astrocytes; OPC, oligodendrocyte progenitor cells; V-SVZ, ventricular-subventricular zone. **c-d**, UMAP plots colored by chromatin accessibility (left) and chromatin velocity (right) of the *Neurod2* promoter region (chr11:98329299-98330151) (**c**) and enhancer region (chr11:98320243-98320844) (**d**). **e**, Dynamics of *Neurod2* in each modality along the pseudotime axis. **f**, Heatmaps of chromatin velocity (left) and chromatin accessibility (right) along pseudotime axis. Peaks with high chromatin velocity similarity are classified into nine clusters. **g-h**, Trends in average chromatin velocity within clusters and gene expression corresponding to TF motifs enriched in the cluster. Motifs enriched among the peaks in cluster C (**g**) and cluster F (**h**) are shown.

One of the unique aspects of mmVelo is its ability to estimate chromatin velocity at a single-peak resolution. We present the estimated accessibility changes in the promoter and enhancer regions of *Neurod2* (Fig. 2c,d). The chromatin velocity estimated using mmVelo accurately captured the changes in accessibility along the trajectory of cell differentiation. While promoter regions only undergo upregulation of accessibility during the early stages of differentiation, enhancer regions also undergo subsequent downregulation. *Neurod2* is a TF belonging to the basic Helix-loop-Helix superfamily, and its transient expression confers a neuronal fate to stem cells (Lin *et al*., 2004, Cherry et al., 2011). These results demonstrated that mmVelo accurately infers the dynamics of chromatin accessibility at the single-peak resolution.

Modelling cell state dynamics on multimodal cell states allows mmVelo to simultaneously estimate the velocity of multiple modalities for each cell, enabling a comparative analysis of the velocities of these modalities. To verify this hypothesis, we investigated the velocity of *Neurod2* along the pseudotime axis for each modality in the excitatory neuron lineage (Fig. 2e). We confirmed a trend in which changes in the enhancer region occurred first, followed by changes in the promoter region, and finally, changes in gene expression. This is consistent with the findings of S. Ma *et al*., 2020, who reported that the regulatory regions of lineage-determining genes were accessible prior to the onset of gene expression. This result suggests that mmVelo possesses the ability to elucidate the temporal dynamics across modalities.

Next, we hypothesized that the rate of change in chromatin accessibility reflects the effects of regulation more directly than chromatin accessibility itself, and that chromatin regions affected by common regulators exhibit synchronous accessibility dynamics. To test this hypothesis, we conducted peak clustering based on chromatin velocity. Nine peak clusters were identified using the Leiden clustering method (Traag, Waltman, and van Eck, 2019). These clusters revealed patterns of change at various stages of development, capturing accessibility patterns that changed synchronously with pseudotime (Fig. 2f).

Similar change patterns observed within each cluster suggest the presence of common regulatory factors. To identify the putative TFs that govern these accessibility changes, we conducted motif enrichment analysis for each peak cluster using pycisTarget. Next, we explored the relationships between the expression patterns of TFs (corresponding to the top-enriched motifs in each cluster) and the average chromatin velocity of the peaks belonging to the cluster (Fig. 2g, h). Notably, *Zbtb2* in cluster F strongly correlated with chromatin velocity and TF expression, suggesting that it plays a role in regulating chromatin accessibility. *Zbtb2* recruits chromatin remodelers and histone chaperones to control differentiation, highlighting its regulatory role in chromatin accessibility (Olivieri *et al*., 2021). These findings demonstrate that mmVelo estimates of temporal changes in chromatin accessibility facilitate the identification of factors that govern the chromatin landscape.

### 2.3 mmVelo reveals the dynamics of transcription factor binding motifs in mouse hair follicle development

Next, we applied mmVelo to mouse skin hair follicle differentiation processes assayed using SHARE-seq (S. Ma *et al*., 2020). The inferred cell state dynamics and chromatin velocity accurately captured the direction of differentiation, as reported in the initial study (Fig. 3a,b). This progression involves the differentiation of transit-amplifying cells into the internal root sheath, medulla, and cuticle/cortex(B. Zhang *et al*., 2016, B. Zhang and Hsu., 2017). Furthermore, benchmark analyses using hair shaft-cuticle/cortex lineage cells demonstrated that mmVelo consistently estimated chromatin velocity and RNA velocity with higher accuracy than existing models, including MultiVelo (C. Li *et al*., 2023) and scVelo (Bergen *et al*., 2020) (Fig. S3j-m).

**Figure 3:**
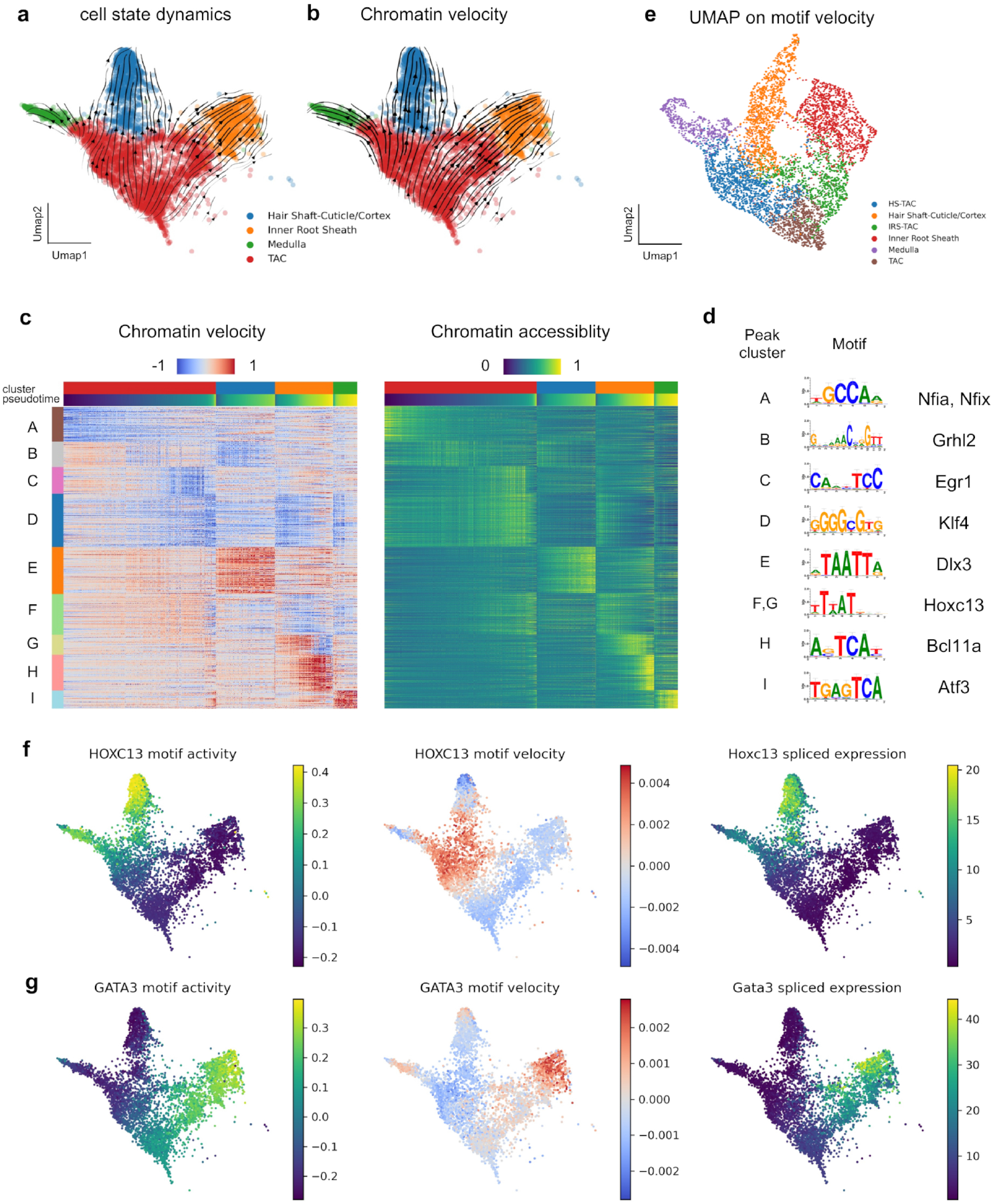
mmVelo reveals the dynamics of TF binding motifs in mouse hair follicle development. **a-b**, UMAP coordinates with stream plots of cell state dynamics (**a**) and chromatin velocity (**b**) from mmVelo. TAC, transit-amplifying cells. **c**, Heatmaps of chromatin velocity (left) and chromatin accessibility (right) along pseudotime axis. Peaks with high similarity in chromatin velocity are classified into nine clusters. **d**, Top enriched TF motifs in each peak cluster. **e**, UMAP representations of the motif velocity. **f-g**, UMAP plots colored by motif activity (left), motif velocity (middle), and spliced mRNA expression (right) of *Hoxc13* (**f**) and *Gata3* (**g**).

To identify TFs that regulate chromatin accessibility, we performed Leiden clustering on chromatin velocity-based peaks estimated using mmVelo. Nine distinct clusters were identified (Fig. 3c). Using pycisTarget (Bravo González-Blas *et al*., 2023), we conducted a motif enrichment analysis for each cluster of peaks and confirmed the enrichment of TF binding motifs (Fig. 3d). Among the enriched motifs, *Nfix*, an NFI TF involved in maintaining stem cell identity by preserving chromatin accessibility (Adam *et al*., 2020), and *Hoxc13*, which is essential for proper hair follicle differentiation (Jave-Suarez *et al*., 2002) were identified. These findings suggest that the chromatin velocity inferred by mmVelo facilitates the elucidation of dynamic regulation and concurrent changes in chromatin accessibility.

To further elucidate the dynamic regulation of TF binding motifs associated with differentiation, we computed the accessibility dynamics of these motifs, which we refer to as motif velocity (Methods). Dimensional reduction of the motif velocity using UMAP recapitulated the continuous transitional changes in cell populations associated with hair follicle differentiation (Fig. 3e), indicating that the estimated motif velocity captured the biological variations in cell states. In particular, the motif velocities of the TFs *Hoxc13* and *Gata3*, which are essential for differentiation into the hair shaft and internal root sheath lineages (Kaufman *et al*., 2003), respectively, accurately captured the changes in motif activity that occur during differentiation (Fig. 3f, g). Furthermore, these changes were concurrent with the gene expression patterns, illustrating the synchronized regulation of the cistrome with gene expression during the differentiation process. These findings demonstrate that in addition to estimating the dynamics of chromatin accessibility, the mmVelo framework enables the estimation of cistrome dynamics.

### 2.4 Chromatin velocity enables inference of putative regulatory TFs for each regulatory region

In the previous section, we showed that considering chromatin velocity alongside TF gene expression allows for the exploration of factors regulating chromatin accessibility. Based on this concept, we hypothesized that chromatin velocity utilization would enable the identification of TFs involved in regulating individual peaks of chromatin accessibility. To demonstrate this, we estimated the regulatory relationships between TFs and chromatin accessibility peaks.

We inferred the regulatory relationships between TFs and chromatin accessibility peaks using a three-step process (Fig. 4a, Methods), grounded in the SCENIC+ framework (Bravo González-Blas *et al*., 2023). Initially, motif enrichment analysis for each cluster of peaks was conducted to identify the candidate TFs involved in regulation. Subsequently, we quantified the importance scores of candidate regulatory TFs in the regression task of chromatin velocity based on their mRNA expression. Finally, the regulatory relationships between TFs and chromatin accessibility peaks were estimated based on importance scores. In our experiment, we estimated the regulatory relationships between 101,644 TF-peak pairs.

**Figure 4:**
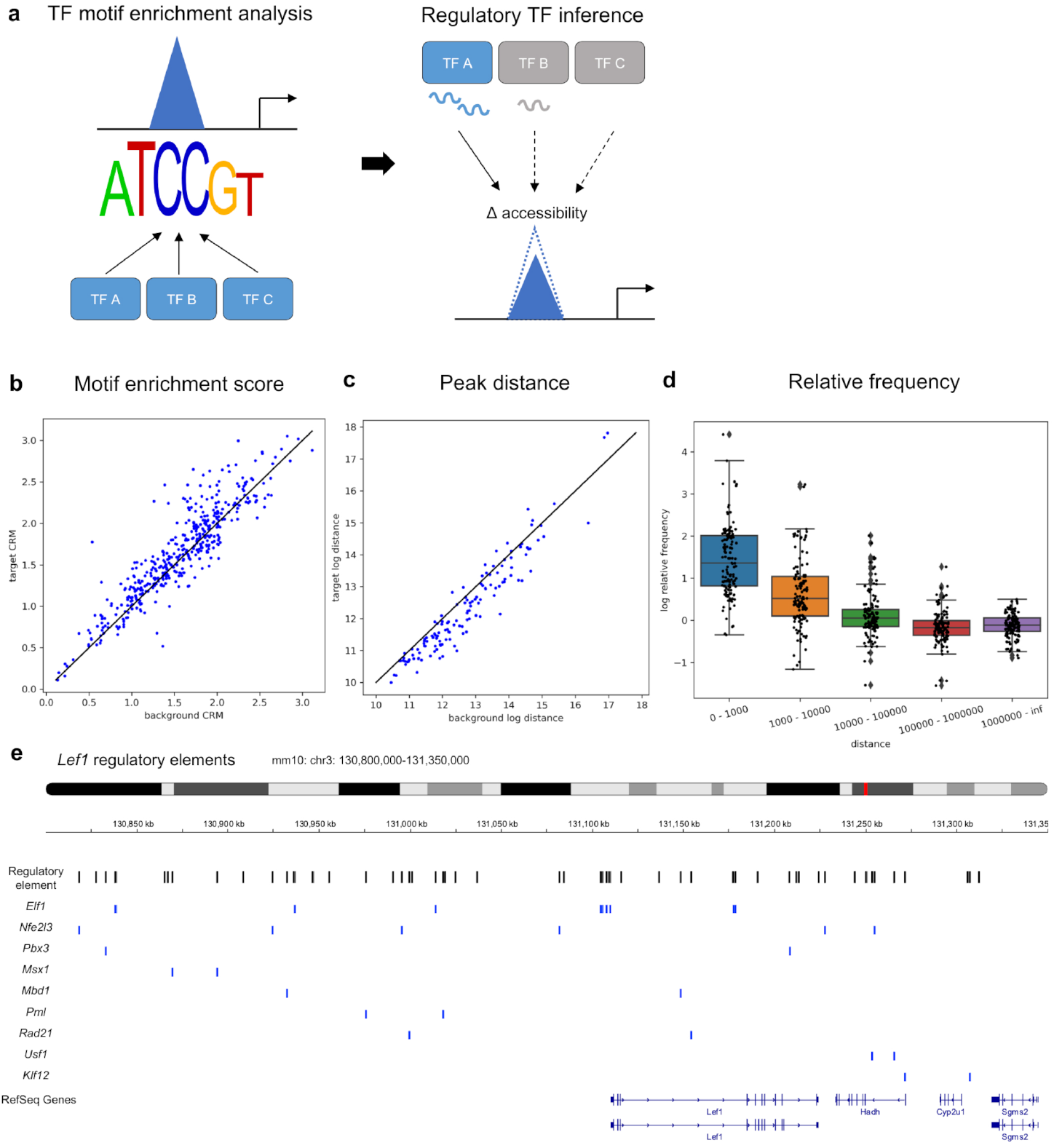
Chromatin accessibility dynamics enables inference of putative regulatory TFs for each regulatory region. **a**, Illustration of regulatory relationship inference between TFs and chromatin accessibility peaks. **b**, Scatter plot representing the CRM scores of TF motifs for TF-targeted peaks (y-axis) and non-targeted peaks within the same cluster (x-axis). Each point corresponds to a TF motif. **c**, Scatter plot representing the mean genomic distances (in base pairs) between TF-targeted peaks (x-axis) and non-targeted peaks (y-axis) on the same chromosome. **d**, Distributions of log-relative frequencies of distances between peaks among targeted and non-targeted peaks. **e**,The inferred regulatory relationships between TF and peaks within the *Lef1* gene regulatory elements (chr3: 130,800,000–131,350,000).

To validate the accuracy of the inferred regulatory relationships, we compared the degree of motif enrichment between TF-regulated and non-regulated peak sets within the same peak clusters. The average likelihood of occurrence of relevant motifs (CRM scores; Cis regulatory modules scores) in the TF-regulated peak sets was higher than that in the non-regulated peak sets (Fig. 4b) (p<0.05, Wilcoxon rank-sum test). This suggests that the extracted regulated peaks contained sequences that were more likely bound by the respective TFs. These results indirectly corroborated the accuracy of the estimated regulatory relationships.

Topologically Associated Domains (TADs) are regions within the genome characterized by self-interacting sequences. The DNA sequences within each TAD interact with each other at a higher frequency than those outside the domain (Pombo and Dillon *et al*., 2015 and Tena and Santos-Pereira, 2021). Inspired by this concept, we hypothesized that TF-regulated chromatin accessibility tends to be concentrated in regions that are genomically proximate. To test this hypothesis, we compared the distances between TF-regulated peaks to the distances between random peaks within the same chromosome. Consistent with our hypothesis, the distances between the regulated peaks were shorter than those between the random peaks (Fig. 4c) (p<0.01, Wilcoxon rank-sum test). To determine the extent to which each TF’s regulated peak is concentrated within a certain distance range, we compared the distribution of the genomic distance between the peaks regulated by a common TF and those between randomly selected peaks. The results showed the concentration of TF-regulated peaks within a range of up to 100 kb (Fig. 4d). These results suggest that the regulation of chromatin accessibility by TFs is concentrated within localized regions of the chromosome.

As an example of the inferred regulatory relationships, the interactions between TFs and peaks in the *Lef1* regulatory elements are shown in (Fig. 4e). For each peak, we depicted the TF predicted to have the most significant positive regulatory effect. *Elf1* and *Nfe2l3* were identified as the TFs that regulate the largest number of regulatory sequences. Notably, *Elf1* was predominantly localized near the coding regions of *Lef1*, consistent with its role as a transcription elongation factor that binds to these regions (Prather *et al*., 2005). These results demonstrate that chromatin velocity utilization facilitates regulatory relationship inference.

### 2.5 mmVelo estimates velocity in missing modalities

Although single-cell multiomics technologies are advancing, these methods are still expensive and less universally applicable than single-modality measurements such as scRNA-seq and scATAC-seq. Therefore, the ability to predict missing modalities from single-modality data provides more profound insights into biological systems in a cost-effective manner. Although several methods can estimate profiles in missing modalities (Minoura *et al*., 2021, Ashuach et al., 2023), no current method can estimate the dynamics of missing modalities. To address this, we developed a method to jointly estimate RNA and chromatin velocities from singleome data via scRNA-seq, scATAC-seq, and Multiome data integration into the mmVelo framework (Fig. 5a, Methods).

**Figure 5:**
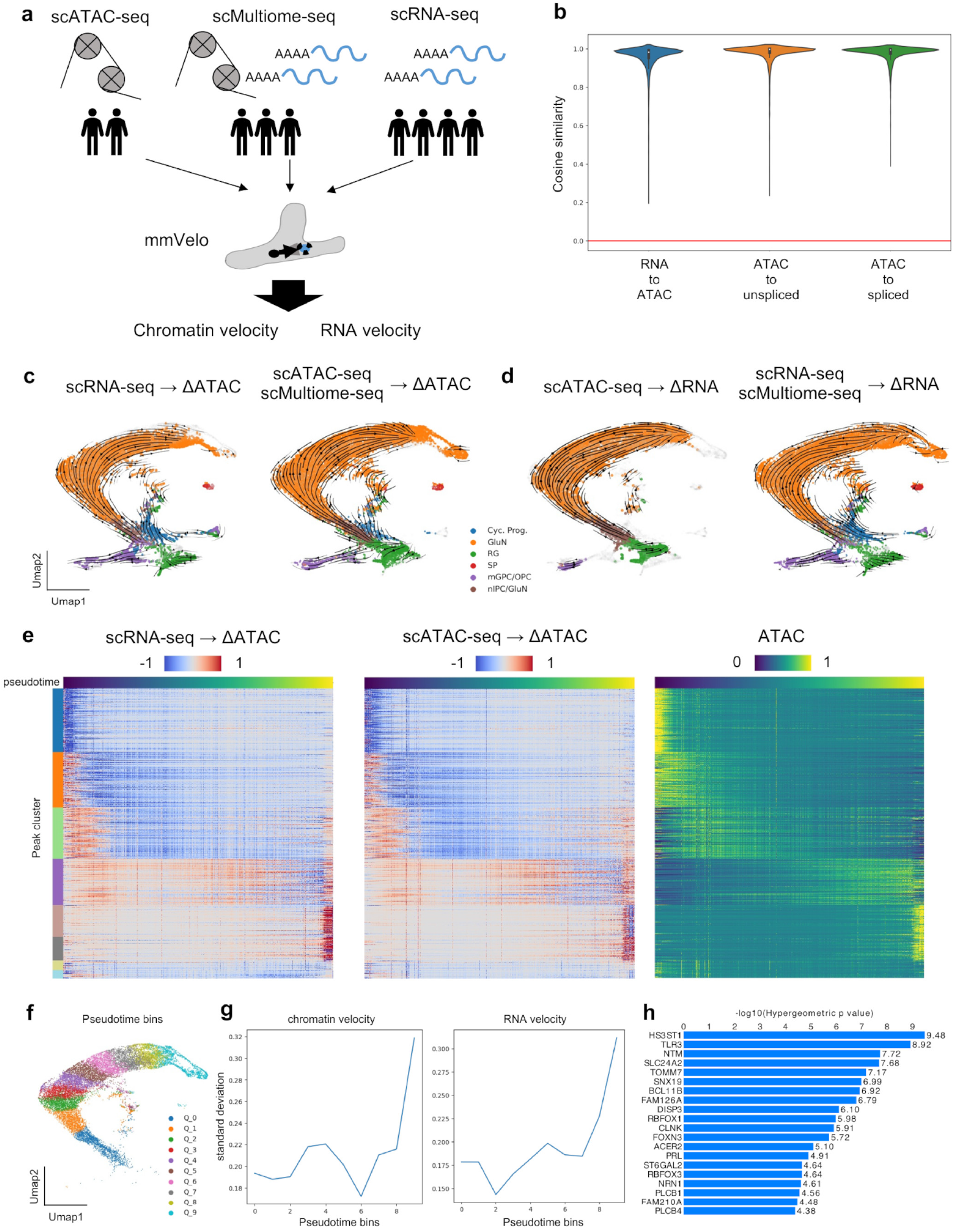
mmVelo estimates velocity in missing modalities. **a**, Diagram of dynamics estimation experiment in missing modality. **b**, Violin plot showing the average cosine similarity between the velocity estimated in missing modality from single-modality data and the velocity in the 10 nearest neighbor cells from multiome data. **c**, UMAP coordinates with stream plot of chromatin velocity inferred from scRNA-seq (left) and from Multiome and scATAC-seq (right). Cyc.Prog., cycling progenitors; GluN, glutamatergic neuron; RG; radial glia; SP, subplate; mGPC/OPC, multipotent glial progenitor cell/ oligodendrocyte progenitor cell; nIPC, neuronal intermediate progenitor cell. **d**, UMAP coordinates with stream plot of RNA velocity inferred from scATAC-seq (left) and from Multiome and scRNA-seq (right). **e**, Heatmaps of chromatin velocity inferred from scRNA-seq (left), scATAC-seq (middle), and chromatin accessibility (right) along pseudotime axis in GluN lineage. **f**, UMAP coordinates of excitatory neuron lineage cells colored by pseudotime bins. **g**, Trends in the average inter-sample variability of chromatin velocity (left) and RNA velocity (right) within each pseudotime bin. **h**, Genes enriched in the top 10 percent of peaks showing high inter-sample variability in the Q4 bin, identified by hypergeometric testing.

As a proof-of-concept, we generated a simulated dataset, with a missing modality, from a 10x Multiome dataset of human cortical development (Trevino *et al*., 2021) and evaluated the estimated velocity of the missing modality. Multiomics data derived from three samples were manipulated such that one sample had an artificially induced missing RNA modality and the other had a missing ATAC modality. We applied mmVelo to this simulated dataset to estimate the velocity of the missing modality. The velocities inferred for the missing modality by mmVelo were consistent with the mean velocities of 10 neighboring multiome cells (Fig. 5b), demonstrating that the model is capable of accurately estimating the dynamics of missing modalities.

Next, we applied mmVelo to a dataset comprising scRNA-seq, scATAC-seq, and 10x Multiome data obtained from the human cortical development process, all measured at the same post-conceptional week (PCW21). Vector fields based on RNA velocity estimated from scATAC-seq and chromatin velocity estimated from scRNA-seq correctly captured the known biological directionality of differentiation (Fig. 5c,d). Furthermore, the dynamics estimated for the missing modalities in the excitatory neuron lineage captured the observed temporal changes in accessibility and gene expression (Fig. 5e). These results demonstrate that mmVelo accurately predicts temporal changes in missing modalities.

The incorporation of singleome data facilitates an increase in the number of samples simultaneously analyzed. In our experiment, integrating multiome data from three individuals with scRNA-seq and/or scATAC-seq data from four additional individuals enabled statistical inferences using a total of seven specimens. Leveraging this, we analyzed the inter-individual variability in dynamics, focusing specifically on the lineage of excitatory neurons, to identify the stage of cell differentiation at which the variability in dynamics becomes pronounced. To quantify the variability in chromatin and RNA velocity among individuals, we divided the cells into ten bins based on pseudotime and computed the variance in the mean velocity for each sample within each bin (Fig. 5f, Methods). Large velocity fluctuations among the samples primarily occurred in chromatin accessibility, followed by changes in gene expression (Fig. 5g). To investigate the functional significance of the top fluctuating peaks, hypergeometric tests were conducted on these peak-associated genes using GREAT (McLean *et al*., 2010). Among the top fluctuating genes, several were associated with the cerebral cortex development or with neurological disorders (Fig. 5h) (Witoelar *et al*., 2018, X. Zhang et al., 2023, L. Ma et al., 2020, Lennon et al., 2017, Miyamoto et al., 2014, Konířová et al., 2017, Fogel et al., 2012). This suggests that certain regulatory mechanisms in neuronal differentiation exhibit significant inter-individual variability, potentially reflecting underlying biological or environmental differences. These results demonstrate the utility of integrating single-modality data for velocity analysis to gain biological insights, highlighting the potential of mmVelo to enhance our understanding of the complex dynamics of cellular processes among different individuals.

## 3 Discussion

RNA velocity analysis is a powerful method for uncovering the dynamics of gene expression within cells. However, existing approaches are limited to RNA modalities, and estimating velocity for other modalities, such as chromatin accessibility, remains a major challenge. To address this gap, we introduce mmVelo, a framework based on a mixture of experts variational autoencoder designed to infer multimodal cell states and their dynamics across multiple modalities.

A key innovation of mmVelo is its ability to model chromatin velocity at a single-peak level, which enables the detailed analyses of time lags between regulatory regions of the same gene, TF motif dynamics, and regulatory relationships with a single-peak resolution. Our experiments revealed how the promoter and enhancer regions of *Neurod2* progressively become accessible over time. By analyzing individual peak dynamics, we inferred the behavior of aggregated cistromes. This was accomplished by extending the motif deviation score from chromVAR to calculate motif velocity (Schep *et al*., 2017), allowing us to quantify the rate of change in lineage-determining TF motifs critical for mouse hair follicle differentiation. Additionally, we extended the SCENIC+ framework to regress chromatin velocity from TF mRNA expression (Bravo González-Blas *et al*., 2023), enabling the identification of transcription factors that regulate peak accessibility. These advancements surpass existing methods, such as MultiVelo (C. Li *et al*., 2023), which aggregate chromatin accessibility at the gene level, providing high-resolution dynamic analyses and deeper insights into regulatory mechanisms across multiple modalities.

In addition, mmVelo’s multimodal representation learning, built on a mixture of experts variational autoencoder framework, allows for the prediction of velocities in missing modalities. For example, using a human cortical development dataset, we demonstrated that both RNA velocity and chromatin velocity can be inferred from scATAC-seq data. While existing models such as scMM (Minoura *et al*., 2021) and MultiVI (Ashuach *et al*., 2023) predict missing modality profiles, they do not estimate the dynamics of these modalities. By applying mmVelo to large single-modality datasets and bridging them with multiomics datasets, researchers can expand the scope of existing data analyses, uncovering novel insights into cellular dynamics and their regulation.

A promising future direction is to integrate mmVelo-based velocity estimation with various mathematical models beyond RNA velocity. For instance, models such as MultiVelo (C. Li *et al*., 2023), which is based on chromatin-gene regulatory relationships, or scKINETICS (Burdziak *et al*., 2023), which is based on gene regulatory networks, can be employed. This integration could offer a more detailed exploration of regulatory relationships within and between modalities and enable dynamic estimation in a broader range of systems, including those in which RNA velocity applications are challenging (Bergen *et al*., 2021). Additionally, combining this approach with fate mapping tools, such as CellRank (Lange *et al*., 2022, Weiler et al., 2024), could help to identify regulatory regions that contribute to cell lineage decisions.

Another important direction is to extend mmVelo to incorporate condition-specific dynamics and identify the contributing factors. Although our model currently assumes that all cells follow the same dynamics, it is plausible that different dynamics exist under specific conditions, such as under different diseases. While studies have been conducted on condition-specific dynamics estimation at the gene expression level (Kojima *et al*., 2024), research extending such estimations across multiple modalities is yet to be conducted. By elucidating the dynamics across modalities and their contributing factors under specific conditions, we can advance our understanding of specific disease mechanisms and drug responses across various molecular layers.

In summary, mmVelo enables the estimation of the dynamics of each modality from single-cell multiomics data. We anticipate that mmVelo will maximize the potential of multiomics data, providing insights into the regulatory relationships across various molecular layers in diverse biological systems, including development, differentiation, and disease.

## 4 Limitations of the Study

mmVelo estimation of cell state dynamics is based on RNA velocity; therefore, it cannot be applied to biological systems where RNA velocity-based estimation of the directionality of cell state changes is difficult (Bergen *et al*., 2021). Nevertheless, it is crucial to note that the dynamics estimation method proposed in this study can be applied to systems that previously challenged traditional RNA velocity methods (e.g., see the application of RNA velocity to hair follicle differentiation in S. Ma *et al*., 2020). Another limitation of our model is the use of smoothed profiles to address the sparsity of observations (Gorin *et al*., 2022, Zheng et al., 2023), along with the assumption of splicing kinetics based on a single kinetic rate (Bergen *et al*., 2021). Expanding the model to include count modeling (Gorin and Pachter, 2022, Lederer *et al*., 2024) and estimating the dynamics based on variable kinetic rates (S. Li *et al*., 2023, Mizukoshi et al., 2024) are future research directions.

## 5 Methods

### 5.1 Overview of the mmVelo algorithm

mmVelo estimates chromatin accessibility dynamics in three stages. Initially, cell states are inferred by leveraging information from all modalities of multiomics data. Subsequently, cell state dynamics are estimated using cell-wise transcriptional profiles and splicing kinetics. Finally, chromatin accessibility dynamics are estimated by mapping cell state dynamics onto the space of chromatin accessibility. Detailed explanations of each of these processes are provided in the following sections.

### 5.2 Cell state inference using mixture-of-experts multimodal variational autoencoder (VAE)

First, we infer multimodal cell states using a variational autoencoder (VAE) (Kingma and Welling, 2013, Lopez et al., 2018, and Minoura *et al*., 2021). Let *n* index cells, *p* index chromatin accessibility regions, and *g* index genes. We assumed a generative model in which the observed chromatin accessibility, *a_np_*, along with the gene expression levels of spliced mRNA, *s_ng_*, and unspliced mRNA, *u_ng_*, were generated based on the following formula:

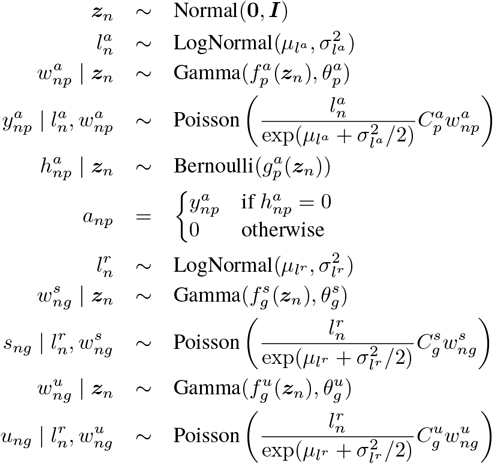

where ***z****_n_* represents the cell state, assumed to follow a normal distribution. The random variables ,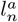 and ,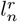 correspond to the scaling factors in scATAC-seq and scRNA-seq, respectively, and were assumed to follow a log-normal distribution. The parameters ,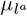 and,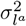, which denote the mean and variance of the log-normal distribution, respectively, were set to the empirical mean and variance of the log-library size (similarly for ,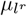 and ,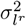 in scRNA-seq). These parameters were tailored to reflect the variability introduced by the differing library sizes among cells. Furthermore ,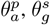, and ,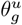 represent the inverse dispersion parameters for each modality, set for each genomic region and gene and optimized during the training process.,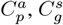, and ,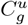 represent the average accessibility scale and average expression scale for peak *p* and gene *g*, respectively. These values were calculated based on the observed counts as ,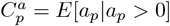 for chromatin accessibility and analogously for spliced and unspliced mRNA.

The functions ,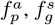, and ,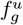 are the decoder neural networks corresponding to each modality. These networks take the cell state ***z****_n_* as the input, and output the mean chromatin accessibility or gene expression for the respective cell. ,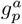 outputs the probability of dropout for a given element, which is a functionality employed to account for the sparsity observed in chromatin accessibility measurements.

Given the difficulty in analytically deriving the posterior distributions of the latent variables,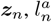, and ,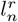, we assume an approximate posterior distribution parameterized by neural networks that take observed values as inputs. This approximation is optimized through variational inference, as follows:

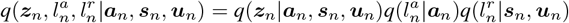

Here, the variational posterior distribution of the cell state is represented as a mixture of experts from the variational posterior distributions of each modality (Shi *et al*., 2019, Minoura et al., 2021), as

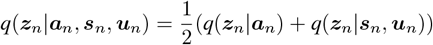

For the variational distribution of the cell state and scaling factor in each modality, we assumed a normal and a log-normal distribution, respectively. The means and variances of these distributions are provided by encoder neural networks that use each modality as an input. Since ,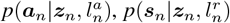 and ,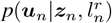 have closed-form probability density functions, the latent variables ,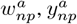 and ,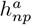 for scATAC-seq, ,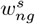 and ,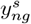 for spliced mRNA, as well as 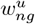 and 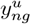 for unspliced mRNA, can be integrated out (Lopez *et al*., 2018). Specifically, ,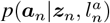 follows a zero-inflated negative binomial distribution with a mean of ,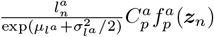, a peak-specific dispersion,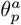, and a zero-inflation probability,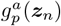, while ,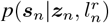 and ,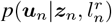 follow negative binomial distributions with means,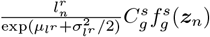 and ,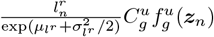 and gene-specific dispersions ,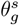 and,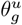, respectively. The model parameters were optimized by maximizing the variational lower bound of the log likelihood:

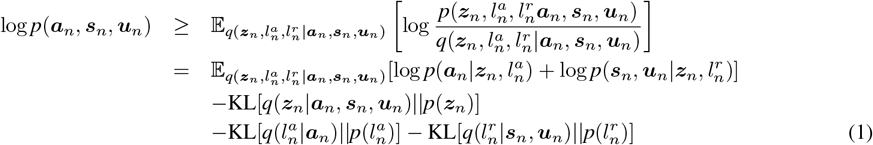

where stratified sampling (Robert and Casella, 2010) is employed for the first term on the right hand side of Equation (1) during training.

### 5.3 Fine-Tuning Decoders for Smoothed Profile Reconstruction

After estimating the cell states through the aforementioned process, mmVelo was fine-tuned to predict the smoothed profiles,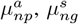, and ,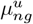 for chromatin accessibility and spliced/unspliced mRNA. The smoothed profiles were calculated based on the neighborhood relationships of the cell states. Specifically, they were determined by taking the average of the size-corrected (to 10,000, Ahlmann-Eltze and Huber, 2023) log1p-normalized counts among the k-nearest neighbor cells. Here, we set *k* = 50. Furthermore, a Gaussian distribution was assumed for the smoothed profiles:

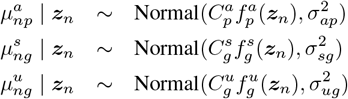

where *σ_ap_, σ_sg_*, and *σ_ug_* are the standard deviations set for each genomic region and gene and are optimized during training process. The model parameters were optimized by maximizing the following term (applied only to the decoder, with all the other parameters of the model held fixed):

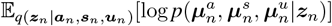

### 5.4 Inference of cell state dynamics

Next, mmVelo inferred cell state dynamics consistent with changes in gene expression described by the RNA velocity equation. This concept was inspired by Nagaharu *et al*., 2022, on which our original formulations were based. At this stage, we assume that the temporal change in the spliced mRNA *d****s****_n_/dt* can be obtained from the following generative model:

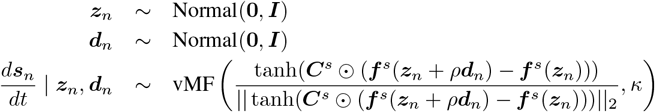

where ⊙ denotes the Hadamard product, *ρ****d****_n_* (*ρ* ≪ 1) represents the transition of the cell state over a short time interval, and ***d****_n_* is assumed to follow a normal distribution. In this study, we set *ρ* = 0.01 to ensure that *ρ****d****_n_* represents a sufficiently small transition in the cell state space. *d****s****_n_/dt* represents the temporal change in the spliced mRNA and is assumed to follow the von Mises-Fisher distribution. The mean of *d****s****_n_/dt* was determined by the difference between the expression profiles before and after a short time interval. The difference in expression profiles was obtained by assuming that the current cellular state is represented by ***z****_n_* and, after a short time interval, by ***z****_n_* + *ρ****d****_n_*, and mapping these states into the gene expression space through the decoder ***f*** ^*s*^ which outputs the average expression profile of spliced mRNA. *κ* is the concentration parameter of the distribution. In this study, we assumed *κ* = 1.

Owing to the difficulty in analytically deriving the posterior distributions of the latent variables ***z****_n_* and ***d****_n_*, we inferred these through variational inference. During this process, we assume that the variational distribution of the cell state transition ***d****_n_* is dependent on the cell state ***z****_n_*, and is determined as follows:

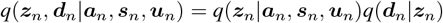

Here, we assumed a normal distribution for the variational posterior distribution of ***d****_n_*, with the mean and variance of this distribution parameterized by an encoder that utilizes the cell state ***z****_n_* as input data. Given the observed values of *d****s****_n_/dt*, optimization was achieved by maximizing the variational lower bound of the log likelihood:

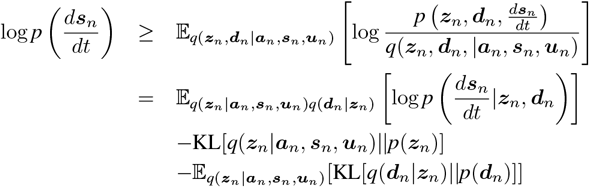

During optimization, we assumed that *d****s****_n_/dt_obs_*, the observed values of *d****s****_n_/dt* were provided based on the assumptions of the splicing kinetics model (La Manno *et al*., 2018) and the output values of the decoders for spliced and unspliced mRNA, ,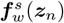 and ,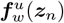, as given by the following equation:

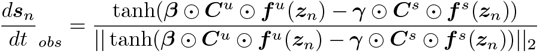

where ***β*** ∈ ℝ ^*g*^ is a vector containing the splicing rates of each gene, and ***γ*** ∈ ℝ^*g*^ is a vector containing the degradation rates of each gene. These were set as learnable parameters and optimized during the training process. The symbol ⊙ denotes the Hadamard product. To prevent genes with large changes in expression levels from being disproportionately emphasized in the inference procedure, the scaling of change in each gene’s expression was performed using the tanh function. During the optimization of the cell state dynamics, all model parameters except the encoder for ***d****_n_* and the splicing kinetics parameters (***β, γ***) were fixed.

### 5.5 Estimation of chromatin velocity

Using the model learned through the above process, we estimated chromatin velocity, the temporal change in chromatin accessibility (ΔATAC). This estimation is achieved by mapping the cellular states during a micro duration to the chromatin accessibility space and calculating their differences, as follows:

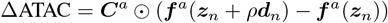

In addition, we estimated the temporal changes in unspliced mRNA (Δunspliced) and spliced mRNA (Δspliced) as follows:

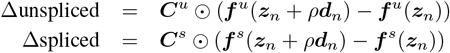

### 5.6 Training procedure

We trained mmVelo on 90% of the data and used 10% as the validation set. By Default, mmVelo was optimized using AdamW (Loshchilov and Hutter, 2017) with a learning rate of 0.0001 for the inference of the cell state and cell state dynamics, and 0.01 for decoder fine-tuning to reconstruct the smoothed profiles. We used a minibatch size of 128 and trained the model for 500 epochs. For each procedure, training was stopped early if there was no improvement in the ELBO on the validation dataset for 30 epochs. As done in a previous study (Minoura *et al*., 2021), for cell state inference, we down-weighted the KL divergence between the cell state variational posterior and its prior for the first 30 epochs.

To ensure that the genes used for estimation had comparable information content in both spliced and unspliced mRNA, we excluded genes with an observed total count ratio of spliced to unspliced mRNA less than 1/50 or greater than 50 during the optimization of cell state dynamics. Additionally, the initial values of the splicing mathematical model parameters ***β*** and ***γ*** were calculated based on the steady-state model of RNA velocity (La Manno *et al*., 2018). Moreover, the squared error of the deviation from the initial values of ***β*** and ***γ*** was included as a regularization term in the optimization objective.

### 5.7 Benchmarking against velocity estimation methods

To benchmark the accuracy of the estimated velocity, we conducted a quantitative comparison of velocity consistency. To this end, we used hair shaft-cuticle/cortex lineage cells during mouse hair follicle development. These cells were chosen because (1) they offer a sufficient number of cells for quantitative evaluation, and (2) their expansion in directions orthogonal to differentiation is relatively modest. Pseudotime for the hair shaft-cuticle/cortex lineage cells was inferred using diffusion pseudotime (Haghverdi *et al*., 2016), and the cell population was subsequently divided into 10 pseudotime bins. Based on these bins, the accuracy of the velocity estimates for each gene or peak was quantified for each method using the velocity consistency score defined by the following equation:

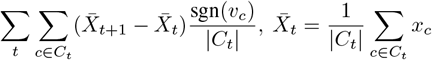

Here, *xc* represents the log-normalized observed count in cell *c*, normalized by dividing the observed count by max(*xc*) to scale the expression. *vc* denotes the estimated velocity in cell *c, t* refers to pseudotime bins, and *Ct* represents the set of cells within pseudotime bin *t*.

For benchmarking, we selected scVelo and MultiVelo (Bergen *et al*., 2020, C. Li *et al*., 2023). To ensure a fair comparison, the neighborhood relationships among cells were computed based on cell states similar to mmVelo, which were estimated using the VAE module of mmVelo. These relationships were then used to smooth the profiles provided as inputs to scVelo and MultiVelo.

### 5.8 Calculation of motif velocity

To estimate the motif velocity, we expanded upon the motif activity concept, as proposed in chromVAR (Schep *et al*., 2017). Let *y_n_* be the motif activity of a motif in cell *n*, and *dy_n_/dt* be the corresponding motif velocity. Initially, the motif activity was calculated in a manner similar to chromVAR based on the following equation:

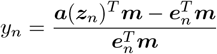

where ***a***(***z****_n_*) represents the reconstructed chromatin accessibility profiles of cell *n*, ***e****_n_* is the expected value of the read count fragments in cell *n*, and ***m*** is a binary vector indicating the presence of a motif at individual peaks, derermined using the motifmatchr package (Schep *et al*., 2017). Specifically, ***e****_n_* is the row vector of ***E***, where *E_ij_* is calculated as *E_ij_* = Σ_*p*_ *a_ip_*(***z***_*i*_) Σ_*n*_ *a_nj_*(***z***_*n*_)*/* Σ_*n*_ Σ_*p*_ *a_np_*(***z***_*n*_). Although chromVAR accounts for technical biases using background peaks, we did not include this correction for this term because we used smoothed profiles and did not observe these biases. Based on this, the motif velocity was calculated using the following formula:

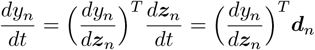

The position-frequency matrix for each TF motif was downloaded from JASPAR2022 (Castro-Mondragon *et al*., 2022), using the getMatrixSet function in Signac (Stuart *et al*., 2021). TF motifs with corresponding TF mRNA expression were used for analysis.

### 5.9 Inferring TF-peak regulatory relationships using chromatin velocity

To identify the TFs that regulate chromatin accessibility, we integrated the SCENIC+ framework (Bravo González-Blas *et al*., 2023) with the estimated chromatin velocity using mmVelo to infer regulatory relationships. Initially, to identify covarying peak clusters, we conducted Leiden clustering of peaks based on chromatin velocity with a resolution of 0.8, employing cosine similarity as a metric for the similarity between peaks. Subsequently, using pycisTarget within the SCENIC+ framework, we conducted a motif search for each peak cluster and designated the corresponding TFs as putative regulators. Finally, we applied GRNBoost2 (Moerman *et al*., 2019) to regress chromatin velocity from the spliced mRNA expression of putative regulator TFs and calculated the feature importance scores for each TF.

To statistically determine the presence of regulatory relationships based on the feature importance scores, we conducted a permutation test. To obtain the empirical distribution of background importance scores, we permuted the TF expression matrix in the direction of cells (while we did not shuffle in the direction of genes to preserve the correlation structure between genes) and applied GRNBoost2; this procedure was repeated five times. Subsequently, adjustments were made using the Benjamini-Hochberg method (Benjamini and Hochberg, 1995), and regulatory relationships were extracted at an FDR of 0.001.

### 5.10 Velocity inference on missing modality

To estimate velocity in the missing modality from single-modality data (scATAC-seq or scRNA-seq), we expanded the mmVelo framework. We used Multiome (snATAC-seq + snRNA-seq), scATAC-seq, and scRNA-seq data as input data. The generative and variational models of this hybrid framework were adopted from the mmVelo model.

For the latent cell state inference, to align the latent variables of both modalities for paired data, we employed stratified sampling optimization with a maximum mean discrepancy (MMD) term. To mitigate batch effects, a one-hot vector representing batch conditions was concatenated to the inputs of both the encoder and decoder. Furthermore, a domain adaptation penalty was applied to the latent cell state space to enhance mixing, following methodologies from scVI, MultiVI and DANN (Lopez *et al*., 2018, Ashuach et al., 2023 and Ganin *et al*., 2015).

When calculating smoothed profiles in the neighborhood to fine-tune the model’s decoder, we defined neighborhood relations using only the data where each modality was present and smoothed the observed counts. To predict the smoothed counts, the values of the one-hot vector representing the batch conditions inputted to the decoder were set to zero.

Finally, the cell state-dependent dynamics for each modality were estimated using all cells, employing the same mmVelo framework.

### 5.11 Preprocessing of data

#### 5.11.1 10× embryonic E18 mouse brain

The filtered accessibility matrix for ATAC-seq and position-sorted RNA alignment (BAM) file of E18 mouse embryonic brain data were downloaded from 10x Genomics website. Unspliced and spliced RNA reads were quantified using the Velocyto run10x command (La Manno *et al*., 2018). Genes with a minimum expression level of fewer than 10 counts were excluded. To focus our analysis on actively differentiating cell populations, we initially excluded interneurons, Cajal-Retzius, and microglia cells. We performed gene and cell filtering, clustering, and annotation based on marker gene expression using the standard Scanpy pipeline (Wolf, Angerer, and Theis, 2018). After filtering out cells that did not actively differentiate, the gene expression count data were normalized using the SCTransform function (Hafemeister and Satija, 2019), implemented in Seurat (note that this normalized expression profile was used only for feature preprocessing and not as model inputs). Following normalization, the top 3,000 highly variable genes were selected. For ATAC-seq data, peaks were filtered based on their proximity to transcription start sites (TSS) and their correlation with gene expression. This peak filtering strategy was inspired by the concept proposed in the SHARE-seq study (S. Ma *et al*., 2020): the LinkPeaks function implemented in Signac (Stuart *et al*., 2021) was applied around TSS regions (±50kb) to identify peaks associated with gene expression. Genes exhibiting a significant correlation (p < 0.05) with at least five peaks were identified as domains of regulatory chromatin (DORC) genes, and these genes were added to the list of RNA-seq genes. Next, the LinkPeaks function was applied within ± 500kb of the TSS, and peaks showing high absolute correlation values (>0.01) with any of the RNA-seq genes were selected as features. The resulting filtered count matrix of gene expression and chromatin accessibility was used as the input for mmVelo.

#### 5.11.2 SHARE-seq mouse skin (hair follicle) data

The quantified ATAC-seq expression matrix, RNA alignment BAM file, and cell annotations of the SHARE-seq mouse skin dataset were downloaded from the Gene Expression Omnibus (GEO). Unspliced and spliced RNA reads were quantified using the Velocyto run command. Hair follicle cells (TAC, IRS, medulla, and hair shaft-cuticle/cortex) were extracted based on cell annotations. Genes with a minimum expression level of fewer than 10 counts were excluded. Gene expression count data were normalized using the SCTransform function implemented in Seurat (note that this normalized expression profile was used only for feature preprocessing). Following normalization, the top 3,000 highly variable genes were selected. For ATAC-seq data, peaks were filtered based on their proximity to transcription start sites (TSS) and their correlation with gene expression. First, the LinkPeaks function implemented in Signac was applied around TSS regions (±50kb) to identify peaks associated with gene expression. Genes exhibiting a significant correlation (p < 0.05) with at least 10 peaks were identified as DORC genes, and those genes were added to the list of RNA-seq genes. Next, the LinkPeaks function was applied within ± 500kb of the TSS, and the top 25,000 peaks showing high absolute correlation values (>0.01) with any of the RNA-seq genes were selected as features. The resulting filtered count matrix of gene expression and chromatin accessibility was used as the input for mmVelo.

#### 5.11.3 Human cerebral cortex

The quantified ATAC-seq expression matrix, quantified unspliced and spliced count matrix, and cell annotations were downloaded from the Gene Expression Omnibus (GEO). We initially excluded interneurons, pericytes, endothelial cells, and microglia cells, because these cells do not actively differentiate. Genes with a minimum expression level of fewer than 10 counts were excluded. Gene expression count data were normalized using the SCTransform function implemented in Seurat (note that this normalized expression profile was only used for feature preprocessing). Following normalization, the top 3,000 highly variable genes were selected. For ATAC-seq data, peaks from both multiome and scATAC-seq data were merged. Combined peaks were created using the reduce function in Signac, followed by re-quantification of peak accessibility using the merge function in Signac. Peaks were then filtered based on their proximity to transcription start sites (TSS) and correlation with gene expression in multiome data. First, the LinkPeaks function implemented in Signac was applied around TSS regions (±50kb) to identify peaks associated with gene expression. Genes exhibiting a significant correlation (p < 0.05) with at least 10 peaks were identified as DORC genes, and these genes were added to the list of RNA-seq genes. Next, the LinkPeaks function was applied within ±500kb of the TSS, and peaks showing high absolute correlation values (>0.01) with any of the RNA-seq genes were selected as features. The resulting filtered count matrix of gene expression and chromatin accessibility was used as the input for mmVelo.

## 7 Acknowledgements

This study was funded by several sources. The Grants-in-Aid for Scientific Research (B) (grant no. 20H04281), Grants-in-Aid for Scientific Research on Innovative Areas on Information Physics of Living Matters (grant no. 22H04839), Grant-in-Aid for Transformative Research Areas (platforms for Advanced Technologies and Research Resources) (grant no. 22H04925), Grant-in-Aid for Transformative Research Areas (A) (grant no. 23H04938), and Grant-in-Aid for Research Activity Start-up (grant no. 20K22839) were provided by the Japan Society for the Promotion of Science (JSPS). Additional support was received from RADDAR-J (grant no. JP22ek0109488), the Project for P-PROMOTE (grant nos. JP22ama221215 and JP22ama221501), Brain/MINDS Health and Diseases (grant no. JP22wm0425007), the Interdisciplinary Cutting-edge Research (grant no. JP23wm0325068), and the Advanced Genome Research and Bioinformatics Study to Facilitate Medical Innovation (GRIFIN) (grant no. JP23tm0424226) from the Japan Agency for Medical Research and Development (AMED). The Moonshot Moonshot R&D program (grant no. JPMJMS2025) and ACT-X program (grant no. JPMJAX20AB) also contributed, through the Japan Science and Technology Agency (JST). Further support was provided by the Medical Research Center Initiative for High Depth Omics and Multilayered Stress Diseases at Institute of Science Tokyo. Supercomputing resources were provided by the Shirokane supercomputer at the Human Genome Center of the University of Tokyo, the TSUBAME3.0 supercomputer at the Tokyo Institute of Technology, and the AI Bridging Cloud Infrastructure (ABCI) at the National Institute of Advanced Industrial Science and Technology (AIST). We would like to thank Editage (www.editage.jp) for English language editing.

## 8 Author contributions

Y.K. and K.M. conceived the idea for this study. N.S. formulated the model and performed the experiments under the supervision of Y.K. and T.S.. S.H., K.A. and H.H. advised on model formulations and downstream analyses. All the authors have read and approved the final manuscript.

## 10 Resource availability

### 10.1 Lead contact

Further information and resource requests should be directed to, and will be fulfilled by, the lead contact, Teppei Shimamura.

### 10.2 Materials availability

No new reagents were generated in this study.

### 10.3 Data and code availability

The 10x embryonic mouse brain dataset was downloaded from the 10x website at https://www.10xgenomics. com/datasets/fresh-embryonic-e-18-mouse-brain-5-k-1-standard-2-0-0. The SHARE-seq mouse skin dataset (S. Ma *et al*., 2020) was downloaded from GEO (GSE140203). The 10x human cortical development dataset was downloaded from GEO (GSE162170).

The mmVelo model was implemented in Python using the PyTorch deep learning library, and the code is available at https://github.com/nomuhyooon/mmVelo. All the original codes will be deposited in Zenodo and will be publicly available for publication.

Any additional information required to reanalyze the data reported in this paper is available from the lead contact upon request.

**Figure S1:**
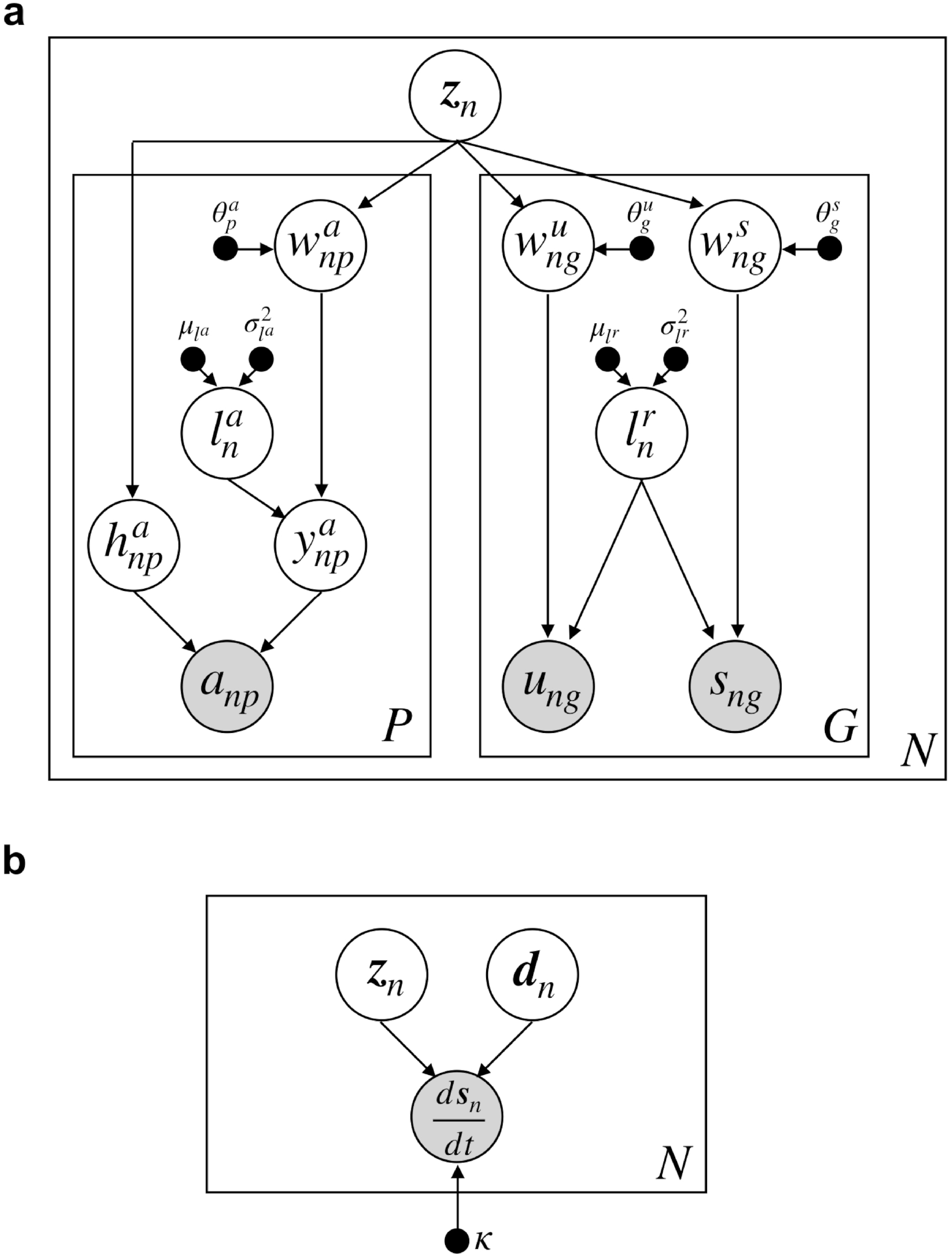
Graphical model of mmVelo. **a**, Graphical model of mmVelo for inferring cell state. **b**, Graphical model of mmVelo for inferring cell state dynamics. Detailed model descriptions and notations are in the Methods section.

**Figure S2:**
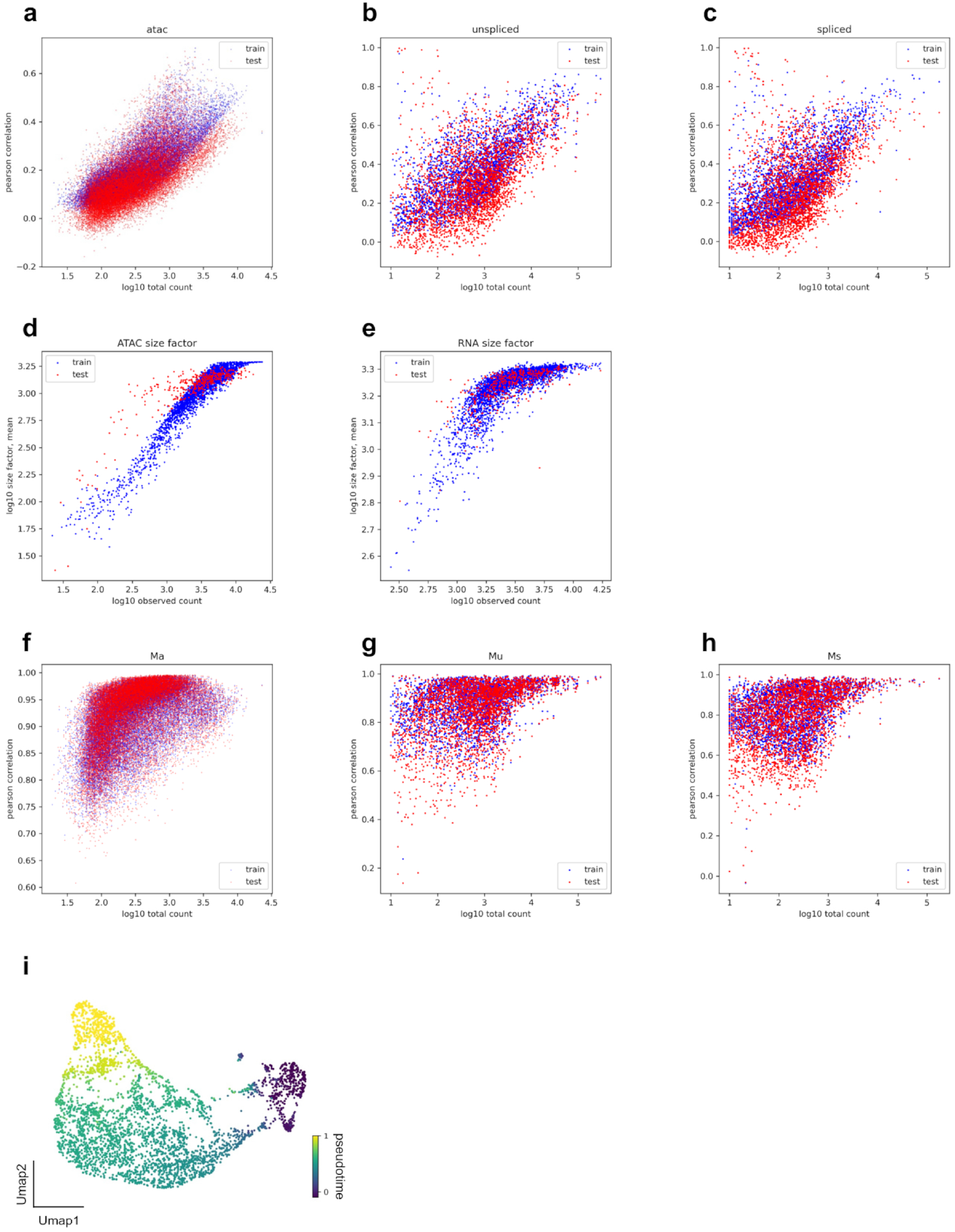
mmVelo model performance in embryonic mouse brain dataset, Related to Figure2. **a-c**, Scatter plot representing the peak-wise/gene-wise correlation between imputed counts in cell state inference and observed counts of ATAC (**a**), unspliced (**b**), and spliced (**c**). **d-e**, Scatter plot representing the cell-wise correlation between inferred size factor in cell state inference and observed library sizes of ATAC (**d**) and RNA (**e**). **f-h**, Scatter plot representing the peak-wise/gene-wise correlation between imputed smoothed profiles and observed moments of ATAC (**f**), unspliced (**g**), and spliced (**h**). **i**, UMAP plots colored by pseudotime. Pseudotime was estimated by diffusion pseudotime, using cell states inferred by mmVelo as inputs.

**Figure S3:**
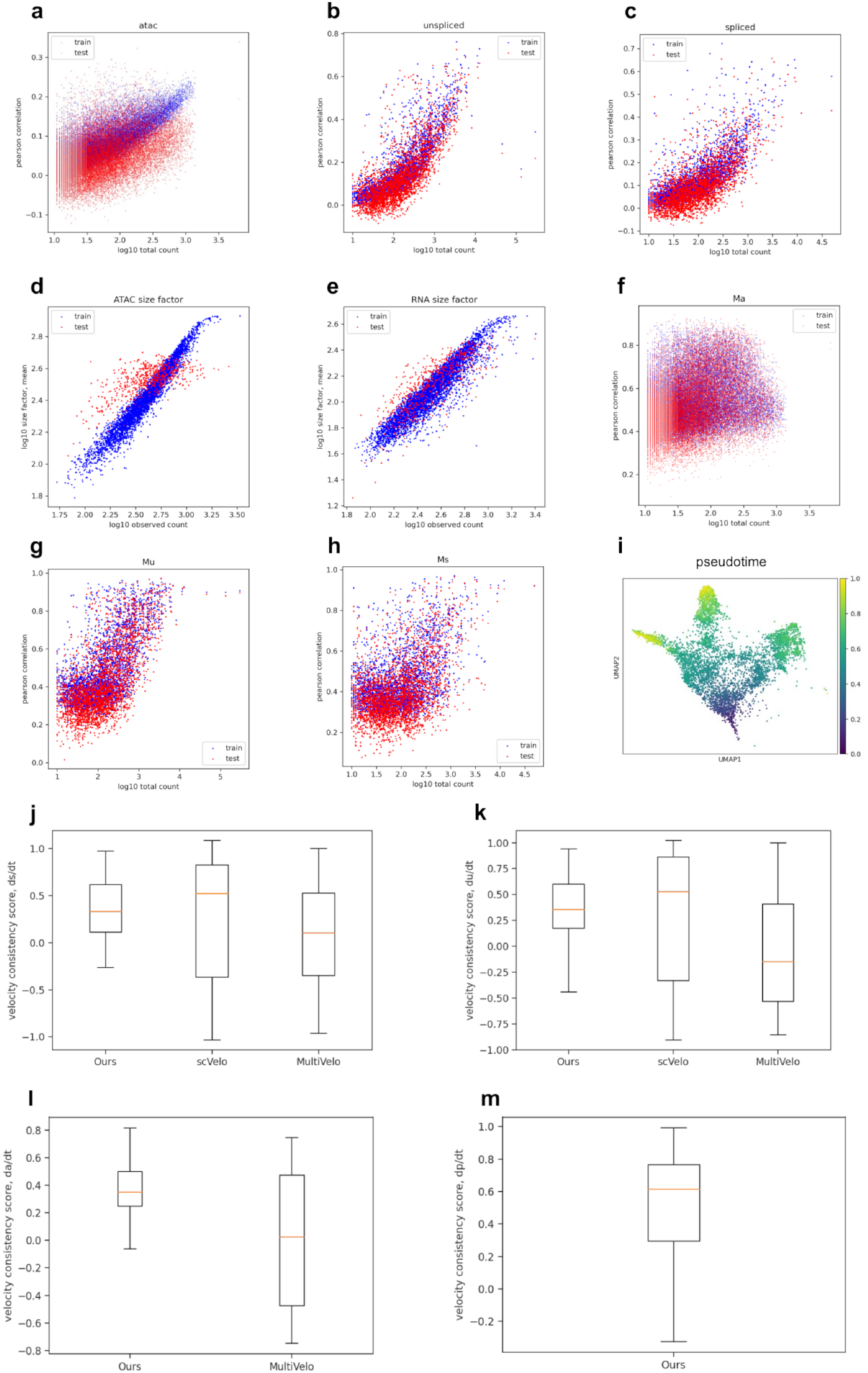
mmVelo model performance in mouse hair follicle development dataset, Related to Figure3. **a-c**, Scatter plot representing the peak-wise/gene-wise correlation between imputed counts in cell state inference and observed counts of ATAC (**a**), unspliced (**b**), and spliced (**c**). **d-e**, Scatter plot representing the cell-wise correlation between inferred size factor in cell state inference and observed library sizes of ATAC (**d**) and RNA (**e**). **f-h**, Scatter plot representing the peak-wise/gene-wise correlation between imputed smoothed profiles and observed moments of ATAC (**f**), unspliced (**g**), and spliced (**h**). **i**, UMAP plots colored by pseudotime. Pseudotime was estimated by diffusion pseudotime, using cell states inferred by mmVelo as inputs. **j-m**, Box plots of velocity consistency score by mmVelo(left), scVelo (middle) and MultiVelo (right) on spliced mRNA velocity (**j**), unspliced mRNA velocity (**k**), gene-wise aggregated chromatin velocity (**l**) and peak-wise chromatin velocity (**m**). In (**l**) scVelo, and in (**m**) scVelo and MultiVelo, are not included because these methods cannot infer the velocity in these modalities. Velocity consistency score was calculated in hair shaft-cuticle/cortex lineage cells. Detailed descriptions about benchmarking are in the Methods section.

**Figure S4:**
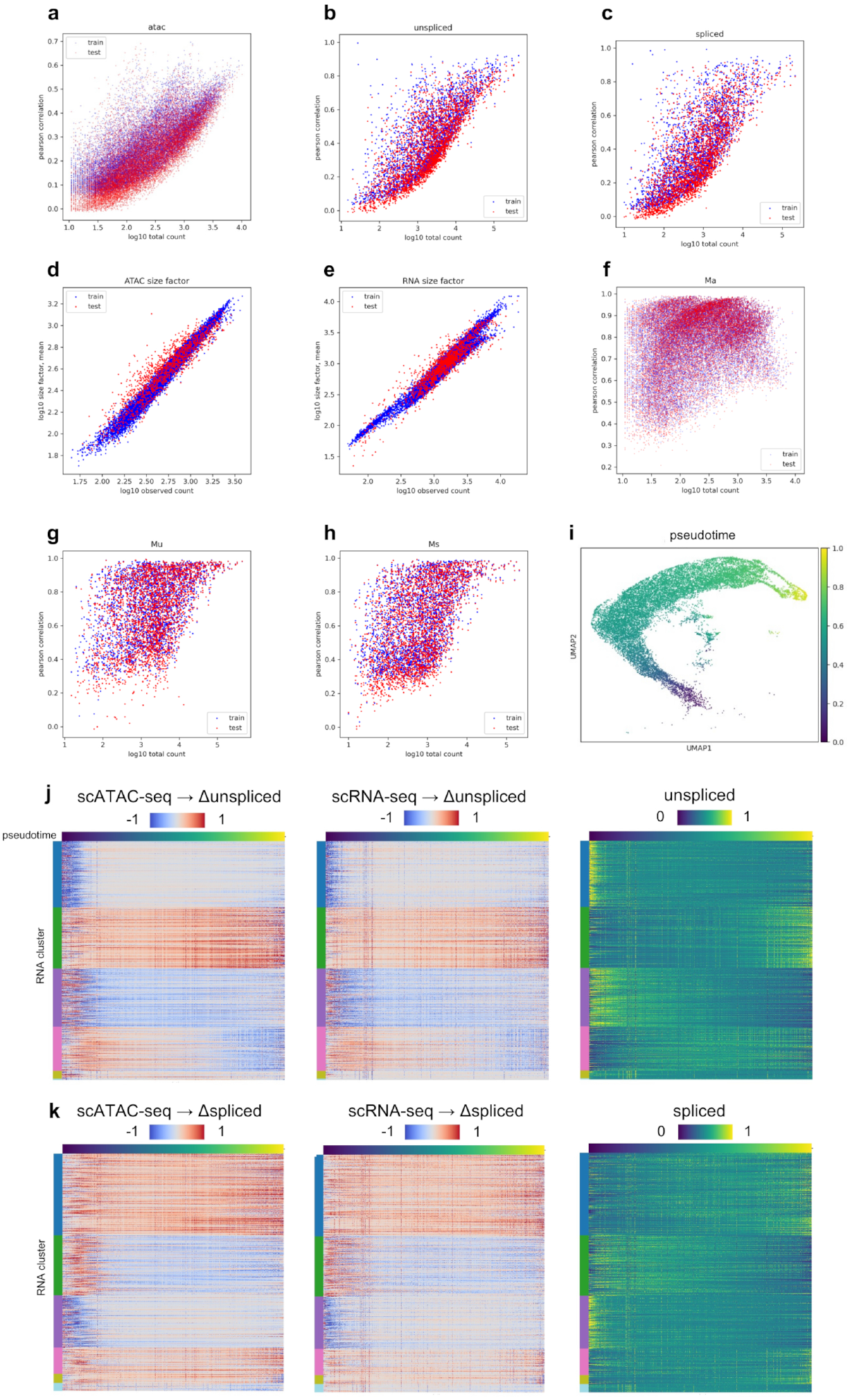
mmVelo model performance in human cortical development dataset, Related to Figure5. **a-c**, Scatter plot representing the peak-wise/gene-wise correlation between imputed counts in cell state inference and observed counts of ATAC (**a**), unspliced (**b**), and spliced (**c**). **d-e**, Scatter plot representing the cell-wise correlation between inferred size factor in cell state inference and observed library sizes of ATAC (**d**) and RNA (**e**). **f-h**, Scatter plot representing the peak-wise/gene-wise correlation between imputed smoothed profiles and observed moments of ATAC (**f**), unspliced (**g**), and spliced (**h**). **i**, UMAP plots of excitatory neuron lineage cells colored by pseudotime. Pseudotime was estimated by diffusion pseudotime, using cell states inferred by mmVelo as inputs. **j-k**, Heatmaps of unspliced mRNA velocity (**j**) and spliced mRNA velocity (**k**) inferred from scATAC-seq (left) and from scRNA-seq (middle), and observed mRNA (right) along pseudotime axis in GluN lineage.

## Reference

Adam, Rene C., Hanseul Yang, Yejing Ge, Nicole R. Infarinato, Shiri Gur-Cohen, Yuxuan Miao, Ping Wang, et al., 2020. “NFI transcription factors provide chromatin access to maintain stem cell identity while preventing unintended lineage fate choices.” Nature Cell Biology 22 (6): 640–50.

Ahlmann-Eltze, Constantin, and Wolfgang Huber. 2023. “Comparison of transformations for single-cell RNA-seq data.” Nature Methods, April. 10.1038/s41592-023-01814-1.

Angermueller, Christof, Stephen J. Clark, Heather J. Lee, Iain C. Macaulay, Mabel J. Teng, Tim Xiaoming Hu, Felix Krueger, et al. 2016. “Parallel single-cell sequencing links transcriptional and epigenetic heterogeneity.” Nature Methods 13 (3): 229–32.

Ashuach, Tal, Mariano I. Gabitto, Rohan V. Koodli, Giuseppe-Antonio Saldi, Michael I. Jordan, and Nir Yosef. 2023. “MultiVI: Deep generative model for the integration of multimodal data.” Nature Methods, June. 10.1038/s41592-023-01909-9.

Bansal, Mukesh, Giusy Della Gatta, and Diego di Bernardo. 2006. “Inference of gene regulatory networks and compound mode of action from time course gene expression profiles.” Bioinformatics 22 (7): 815–22.

Baysoy, Alev, Zhiliang Bai, Rahul Satija, and Rong Fan. 2023. “The technological landscape and applications of single-cell multi-omics.” Nature Reviews. Molecular Cell Biology, June, 1–19.

Benjamini, Y., and Y. Hochberg. 1995. “Controlling the false discovery rate: A practical and powerful approach to multiple testing.” Journal of the Royal Statistical Society Series B-Methodological 57 (1): 289–300.

Bergen, Volker, Marius Lange, Stefan Peidli, F. Alexander Wolf, and Fabian J. Theis. 2020. “Generalizing RNA velocity to transient cell states through dynamical modeling.” Nature Biotechnology 38 (12): 1408–14.

Bergen, Volker, Ruslan A. Soldatov, Peter V. Kharchenko, and Fabian J. Theis. 2021. “RNA Velocity-Current challenges and future perspectives.” Molecular Systems Biology 17 (8): e10282.

Bravo González-Blas, Carmen, Seppe De Winter, Gert Hulselmans, Nikolai Hecker, Irina Matetovici, Valerie Christiaens, Suresh Poovathingal, Jasper Wouters, Sara Aibar, and Stein Aerts. 2023. “SCENIC+: Single-cell multiomic inference of enhancers and gene regulatory networks.” Nature Methods, July. 10.1038/s41592-023-01938-4.

Burdziak, Cassandra, Chujun Julia Zhao, Doron Haviv, Direna Alonso-Curbelo, Scott W. Lowe, and Dana Pe’er. 2023. “scKINETICS: Inference of regulatory velocity with single-cell transcriptomics data.” Bioinformatics 39 (39 Suppl 1): i394–403.

Cannoodt, Robrecht, Wouter Saelens, and Yvan Saeys. 2016. “Computational methods for trajectory inference from single-cell transcriptomics.” European Journal of Immunology 46 (11): 2496–2506.

Cao, Junyue, Darren A. Cusanovich, Vijay Ramani, Delasa Aghamirzaie, Hannah A. Pliner, Andrew J. Hill, Riza M. Daza, et al. 2018. “Joint profiling of chromatin accessibility and gene expression in thousands of single cells.” Science 361 (6409): 1380–85.

Castro-Mondragon, Jaime A., Rafael Riudavets-Puig, Ieva Rauluseviciute, Roza Berhanu Lemma, Laura Turchi, Romain Blanc-Mathieu, Jeremy Lucas, et al. 2022. “JASPAR 2022: The 9th release of the open-access database of transcription factor binding profiles.” Nucleic Acids Research 50 (D1): D165–73.

Chen, Amy F., Benjamin Parks, Arwa S. Kathiria, Benjamin Ober-Reynolds, Jorg J. Goronzy, and William J. Greenleaf. 2022. “NEAT-Seq: Simultaneous profiling of intra-nuclear proteins, chromatin accessibility and gene expression in single cells.” Nature Methods 19 (5): 547–53.

Chen, Song, Blue B. Lake, and Kun Zhang. 2019. “High-throughput sequencing of the transcriptome and chromatin accessibility in the same cell.” Nature Biotechnology 37 (12): 1452–57.

Cherry, Timothy J., Sui Wang, Ingo Bormuth, Markus Schwab, James Olson, and Constance L. Cepko. 2011. “NeuroD factors regulate cell fate and neurite stratification in the developing retina.” The Journal of Neuroscience: The Official Journal of the Society for Neuroscience 31 (20): 7365–79.

Clark, Stephen J., Ricard Argelaguet, Chantriolnt-Andreas Kapourani, Thomas M. Stubbs, Heather J. Lee, Celia Alda-Catalinas, Felix Krueger, et al. 2018. “scNMT-Seq enables joint profiling of chromatin accessibility DNA methylation and transcription in single cells.” Nature Communications 9 (1): 1–9.

Di Bernardo, D., T. S. Gardner, and J. J. Collins. 2004. “Robust identification of large genetic networks.” Pacific Symposium on Biocomputing 486–97.

Fogel, Brent L., Eric Wexler, Amanda Wahnich, Tara Friedrich, Chandran Vijayendran, Fuying Gao, Neelroop Parikshak, Genevieve Konopka, and Daniel H. Geschwind. 2012. “RBFOX1 regulates both splicing and transcriptional networks in human neuronal development.” Human Molecular Genetics 21 (19): 4171–86.

Ganin, Yaroslav, Evgeniya Ustinova, Hana Ajakan, Pascal Germain, Hugo Larochelle, François Laviolette, Mario Marchand, and Victor Lempitsky. 2015. “Domain-adversarial training of neural networks.” arXiv [stat.ML]. arXiv. http://arxiv.org/abs/1505.07818.

Gardner, Timothy S., Diego di Bernardo, David Lorenz, and James J. Collins. 2003. “Inferring genetic networks and identifying compound mode of action via expression profiling.” Science (New York, N.Y.) 301 (5629): 102–5.

Gayoso, Adam, Zoë Steier, Romain Lopez, Jeffrey Regier, Kristopher L. Nazor, Aaron Streets, and Nir Yosef. 2021. “Joint probabilistic modeling of single-cell multi-omic data with totalVI.” Nature Methods 18 (3): 272–82.

Gorin, Gennady, Meichen Fang, Tara Chari, and Lior Pachter. 2022. “RNA velocity unraveled.” PLoS Computational Biology 18 (9): e1010492.

Gorin, Gennady, and Lior Pachter. 2022. “Modeling bursty transcription and splicing with the chemical master equation.” Biophysical Journal 121 (6): 1056–69.

Hafemeister, Christoph, and Rahul Satija. 2019. “Normalization and variance stabilization of single-cell RNA-Seq data using regularized negative binomial regression.” Genome Biology 20 (1): 296.

Haghverdi, Laleh, Maren Büttner, F. Alexander Wolf, Florian Buettner, and Fabian J. Theis. 2016. “Diffusion pseudotime robustly reconstructs lineage branching.” Nature Methods 13 (10): 845–48.

Hansen, David V., Jan H. Lui, Philip R. L. Parker, and Arnold R. Kriegstein. 2010. “Neurogenic radial glia in the outer subventricular zone of human neocortex.” Nature 464 (7288): 554–61.

Hou, Yu, Huahu Guo, Chen Cao, Xianlong Li, Boqiang Hu, Ping Zhu, Xinglong Wu, et al. 2016. “Single-cell triple omics sequencing reveals genetic, epigenetic, and transcriptomic heterogeneity in hepatocellular carcinomas.” Cell Research 26 (3): 304–19.

Hu, Youjin, Kevin Huang, Qin An, Guizhen Du, Ganlu Hu, Jinfeng Xue, Xianmin Zhu, Cun-Yu Wang, Zhigang Xue, and Guoping Fan. 2016. “Simultaneous profiling of transcriptome and DNA methylome from a single cell.” Genome Biology 17 (May):88.

Jave-Suarez, Luis Felipe, Hermelita Winter, Lutz Langbein, Michael A. Rogers, and Jürgen Schweizer. 2002. “HOXC13 is involved in the regulation of human hair keratin gene expression.” The Journal of Biological Chemistry 277 (5): 3718–26.

Kaufman, Charles K., Ping Zhou, H. Amalia Pasolli, Michael Rendl, Diana Bolotin, Kim-Chew Lim, Xing Dai, Maria-Luisa Alegre, and Elaine Fuchs. 2003. “GATA-3: An unexpected regulator of cell lineage determination in skin.” Genes & Development 17 (17): 2108–22.

Kingma, Diederik P., and Max Welling. 2013. “Auto-encoding variational bayes.” arXiv [stat.ML]. arXiv. http://arxiv.org/abs/1312.6114.

Kojima, Yasuhiro, Yuko Arioka, Haruka Hirose, Shuto Hayashi, Yusuke Mizuno, Keiki Nagaharu, Hiroki Okumura, et al. 2024. “Inferring extrinsic factor-dependent single-cell transcriptome dynamics using a deep generative model.” bioRxiv. 10.1101/2024.04.01.587302.

Konířová, Jana, Jana Oltová, Alicia Corlett, Justyna Kopycińska, Michal Kolář, Petr Bartůněk, and Martina Zíková. 2017. “Modulated DISP3/PTCHD2 expression influences neural stem cell fate decisions.” Scientific Reports 7 (1): 41597.

La Manno Gioele, Ruslan Soldatov, Amit Zeisel, Emelie Braun, Hannah Hochgerner, Viktor Petukhov, Katja Lidschreiber, et al. 2018. “RNA velocity of single cells.” Nature 560 (7719): 494–98.

Lange, Marius, Volker Bergen, Michal Klein, Manu Setty, Bernhard Reuter, Mostafa Bakhti, Heiko Lickert, et al. 2022. “CellRank for directed single-cell fate mapping.” Nature Methods 19 (2): 159–70.

Lederer, Alex R., Maxine Leonardi, Lorenzo Talamanca, Daniil M. Bobrovskiy, Antonio Herrera, Colas Droin, Irina Khven, et al. 2024. “Statistical inference with a manifold-constrained RNA velocity model uncovers cell cycle speed modulations.” Nature Methods, October, 1–16.

Lennon, Matthew J., Simon P. Jones, Michael D. Lovelace, Gilles J. Guillemin, and Bruce J. Brew. 2017. “Bcl11b-A critical neurodevelopmental transcription factor-roles in health and disease.” Frontiers in Cellular Neuroscience 11 (March):89.

Li, Chen, Maria C. Virgilio, Kathleen L. Collins, and Joshua D. Welch. 2023. “Multi-omic single-cell velocity models epigenome-transcriptome interactions and improves cell fate prediction.” Nature Biotechnology 41 (3): 387–98.

Lin, Chin-Hsing, Jennifer Stoeck, Ali C. Ravanpay, François Guillemot, Stephen J. Tapscott, and James M. Olson. 2004. “Regulation of neuroD2 expression in mouse brain.” Developmental Biology 265 (1): 234–45.

Li, Shengyu, Pengzhi Zhang, Weiqing Chen, Lingqun Ye, Kristopher W. Brannan, Nhat-Tu Le, Jun-Ichi Abe, John P. Cooke, and Guangyu Wang. 2023. “A relay velocity model infers cell-dependent RNA velocity.” Nature Biotechnology, April. 10.1038/s41587-023-01728-5.

Liu, Longqi, Chuanyu Liu, Andrés Quintero, Liang Wu, Yue Yuan, Mingyue Wang, Mengnan Cheng, et al. 2019. “Deconvolution of single-cell multi-omics layers reveals regulatory heterogeneity.” Nature Communications 10 (1): 1–10.

Lopez, Romain, Jeffrey Regier, Michael B. Cole, Michael I. Jordan, and Nir Yosef. 2018. “Deep generative modeling for single-cell transcriptomics.” Nature Methods 15 (12): 1053–58.

Loshchilov, Ilya, and Frank Hutter. 2017. “Decoupled weight decay regularization.” arXiv [cs.LG]. arXiv. http://arxiv.org/abs/1711.05101.

Luo, Chongyuan, Hanqing Liu, Fangming Xie, Ethan J. Armand, Kimberly Siletti, Trygve E. Bakken, Rongxin Fang, et al. 2022. “Single nucleus multi-omics identifies human cortical cell regulatory genome diversity.” Cell Genomics 2 (3): 100107.

Ma, Baoshan, Mingkun Fang, and Xiangtian Jiao. 2020. “Inference of gene regulatory networks based on nonlinear ordinary differential equations.” Bioinformatics (Oxford, England) 36 (19): 4885–93.

Ma, Liang, Stephen A. Semick, Qiang Chen, Chao Li, Ran Tao, Amanda J. Price, Joo Heon Shin, et al. 2020. “Schizophrenia risk variants influence multiple classes of transcripts of sorting Nexin 19 (SNX19).” Molecular Psychiatry 25 (4): 831–43.

Ma, Sai, Bing Zhang, Lindsay M. LaFave, Andrew S. Earl, Zachary Chiang, Yan Hu, Jiarui Ding, et al. 2020. “Chromatin potential identified by shared single-cell profiling of RNA and chromatin.” Cell 183 (4): 1103–16.e20.

Matsumoto, Hirotaka, Hisanori Kiryu, Chikara Furusawa, Minoru S. H. Ko, Shigeru B. H. Ko, Norio Gouda, Tetsutaro Hayashi, and Itoshi Nikaido. 2017. “SCODE: An efficient regulatory network inference algorithm from single-cell RNA-seq during differentiation.” Bioinformatics (Oxford, England) 33 (15): 2314–21.

Mayer, Simone, Jiadong Chen, Dmitry Velmeshev, Andreas Mayer, Ugomma C. Eze, Aparna Bhaduri, Carlos E. Cunha, et al. 2019. “Multimodal single-cell analysis reveals physiological maturation in the developing human neocortex.” Neuron 102 (1): 143–58.e7.

McLean, Cory Y., Dave Bristor, Michael Hiller, Shoa L. Clarke, Bruce T. Schaar, Craig B. Lowe, Aaron M. Wenger, and Gill Bejerano. 2010. “GREAT improves functional interpretation of cis-regulatory regions.” Nature Biotechnology 28 (5): 495–501.

Merkle, Florian T., Anthony D. Tramontin, José Manuel García-Verdugo, and Arturo Alvarez-Buylla. 2004. “Radial glia give rise to adult neural stem cells in the subventricular zone.” Proceedings of the National Academy of Sciences of the United States of America 101 (50): 17528–32.

Mimitou, Eleni P., Anthony Cheng, Antonino Montalbano, Stephanie Hao, Marlon Stoeckius, Mateusz Legut, Timothy Roush, et al. 2019. “Multiplexed detection of proteins, transcriptomes, clonotypes and CRISPR perturbations in single cells.” Nature Methods 16 (5): 409–12.

Mimitou, Eleni P., Caleb A. Lareau, Kelvin Y. Chen, Andre L. Zorzetto-Fernandes, Yuhan Hao, Yusuke Takeshima, Wendy Luo, et al. 2021. “Scalable, multimodal profiling of chromatin accessibility, gene expression and protein levels in single cells.” Nature Biotechnology 39 (10): 1246–58.

Minoura, Kodai, Ko Abe, Hyunha Nam, Hiroyoshi Nishikawa, and Teppei Shimamura. 2021. “A mixture-of-experts deep generative model for integrated analysis of single-cell multiomics data.” Cell Reports Methods 1 (5): 100071.

Miyamoto, Yuki, Tomohiro Torii, Takahiro Eguchi, Kazuaki Nakamura, Akito Tanoue, and Junji Yamauchi. 2014. “Hypomyelinating leukodystrophy-associated missense mutant of FAM126A/hyccin/DRCTNNB1A aggregates in the endoplasmic reticulum.” Journal of Clinical Neuroscience: Official Journal of the Neurosurgical Society of Australasia 21 (6): 1033–39.

Mizukoshi, Chikara, Yasuhiro Kojima, Satoshi Nomura, Shuto Hayashi, Ko Abe, and Teppei Shimamura. 2024. “DeepKINET: A deep generative model for estimating single-cell RNA splicing and degradation rates.” Genome Biology 25 (1): 229.

Moerman, Thomas, Sara Aibar Santos, Carmen Bravo González-Blas, Jaak Simm, Yves Moreau, Jan Aerts, and Stein Aerts. 2019. “GRNBoost2 and Arboreto: Efficient and scalable inference of gene regulatory networks.” Bioinformatics (Oxford, England) 35 (12): 2159–61.

Nadarajah, Bagirathy, and John G. Parnavelas. 2002. “Modes of neuronal migration in the developing cerebral cortex.” Nature Reviews. Neuroscience 3 (6): 423–32.

Nagaharu, Keiki, Yasuhiro Kojima, Haruka Hirose, Kodai Minoura, Kunihiko Hinohara, Hirohito Minami, Yuki Kageyama, et al. 2022. “A bifurcation concept for B-lymphoid/plasmacytoid dendritic cells with largely fluctuating transcriptome dynamics.” Cell Reports 40 (9): 111260.

Ocone, Andrea, Laleh Haghverdi, Nikola S. Mueller, and Fabian J. Theis. 2015. “Reconstructing gene regulatory dynamics from high-dimensional single-cell snapshot data.” Bioinformatics (Oxford, England) 31 (12): i89–96.

Olivieri, Daniel, Sujani Paramanathan, Anaïs F. Bardet, Daniel Hess, Sébastien A. Smallwood, Ulrich Elling, and Joerg Betschinger. 2021. “The BTB-domain transcription factor ZBTB2 recruits chromatin remodelers and a histone chaperone during the exit from pluripotency.” The Journal of Biological Chemistry 297 (2): 100947.

Pan, Lixia, Wai Lim Ku, Qingsong Tang, Yaqiang Cao, and Keji Zhao. 2022. “scPCOR-Seq enables co-profiling of chromatin occupancy and RNAs in single cells.” Communications Biology 5 (1): 678.

Plongthongkum, Nongluk, Dinh Diep, Song Chen, Blue B. Lake, and Kun Zhang. 2021. “Scalable dual-omics profiling with single-nucleus chromatin accessibility and mRNA expression sequencing 2 (SNARE-seq2).” Nature Protocols 16 (11): 4992–5029.

Pollen, Alex A., Tomasz J. Nowakowski, Jiadong Chen, Hanna Retallack, Carmen Sandoval-Espinosa, Cory R. Nicholas, Joe Shuga, et al. 2015. “Molecular identity of human outer radial glia during cortical development.” Cell 163 (1): 55–67.

Pombo, Ana, and Niall Dillon. 2015. “Three-dimensional genome architecture: Players and mechanisms.” Nature Reviews. Molecular Cell Biology 16 (4): 245–57.

Prather, Donald, Nevan J. Krogan, Andrew Emili, Jack F. Greenblatt, and Fred Winston. 2005. “Identification and characterization of Elf1, a conserved transcription elongation factor in Saccharomyces Cerevisiae.” Molecular and Cellular Biology 25 (22): 10122–35.

Rang, Franka J., Kim L. de Luca, Sandra S. de Vries, Christian Valdes-Quezada, Ellen Boele, Phong D. Nguyen, Isabel Guerreiro, et al. 2022. “Single-Cell profiling of transcriptome and histone modifications with EpiDamID.” Molecular Cell 82 (10): 1956–70.e14.

Robert, Christian, and George Casella. 2010. Monte Carlo statistical methods. Springer Texts in Statistics. New York, NY: Springer.

Rooijers, Koos, Corina M. Markodimitraki, Franka J. Rang, Sandra S. de Vries, Alex Chialastri, Kim L. de Luca, Dylan Mooijman, Siddharth S. Dey, and Jop Kind. 2019. “Simultaneous quantification of protein–DNA contacts and transcriptomes in single cells.” Nature Biotechnology 37 (7): 766–72.

Saelens, Wouter, Robrecht Cannoodt, Helena Todorov, and Yvan Saeys. 2019. “A comparison of single-cell trajectory inference methods.” Nature Biotechnology 37 (5): 547–54.

Schep, Alicia N., Beijing Wu, Jason D. Buenrostro, and William J. Greenleaf. 2017. “chromVAR: Inferring transcription-factor-associated accessibility from single-cell epigenomic data.” Nature Methods 14 (10): 975–78.

Setty, Manu, Vaidotas Kiseliovas, Jacob Levine, Adam Gayoso, Linas Mazutis, and Dana Pe’er. 2019. “Characterization of cell fate probabilities in single-cell data with Palantir.” Nature Biotechnology 37 (4): 451–60.

Setty, Manu, Michelle D. Tadmor, Shlomit Reich-Zeliger, Omer Angel, Tomer Meir Salame, Pooja Kathail, Kristy Choi, Sean Bendall, Nir Friedman, and Dana Pe’er. 2016. “Wishbone identifies bifurcating developmental trajectories from single-cell data.” Nature Biotechnology 34 (6): 637–45.

Shi, Yuge, N. Siddharth, Brooks Paige, and Philip H. S. Torr. 2019. “Variational mixture-of-experts autoencoders for multi-modal deep generative models.” arXiv [stat.ML]. arXiv. http://arxiv.org/abs/1911.03393.

Stuart, Tim, Avi Srivastava, Shaista Madad, Caleb A. Lareau, and Rahul Satija. 2021. “Single-cell chromatin state analysis with Signac.” Nature Methods 18 (11): 1333–41.

Sun, Zhongxing, Yin Tang, Yanjun Zhang, Yuan Fang, Junqi Jia, Weiwu Zeng, and Dong Fang. 2021. “Joint single-cell multiomic analysis in Wnt3a induced asymmetric stem cell division.” Nature Communications 12 (1): 1–19.

Swanson, Elliott, Cara Lord, Julian Reading, Alexander T. Heubeck, Palak C. Genge, Zachary Thomson, Morgan D. A. Weiss, et al. 2021. “Simultaneous trimodal single-cell measurement of transcripts, epitopes, and chromatin accessibility using TEA-Seq.” eLife 10 (April):e63632.

Tedesco, Martina, Francesca Giannese, Dejan Lazarević, Valentina Giansanti, Dalia Rosano, Silvia Monzani, Irene Catalano, et al. 2022. “Chromatin Velocity reveals epigenetic dynamics by single-cell profiling of heterochromatin and euchromatin.” Nature Biotechnology 40 (2): 235–44.

Tena, Juan J., and José M. Santos-Pereira. 2021. “Topologically associating domains and regulatory landscapes in development, evolution and disease.” Frontiers in Cell and Developmental Biology 9 (July):702787.

Traag, V. A., L. Waltman, and N. J. van Eck. 2019. “From Louvain to Leiden: Guaranteeing well-connected communities.” Scientific Reports 9 (1): 5233.

Trapnell, Cole, Davide Cacchiarelli, Jonna Grimsby, Prapti Pokharel, Shuqiang Li, Michael Morse, Niall J. Lennon, Kenneth J. Livak, Tarjei S. Mikkelsen, and John L. Rinn. 2014. “The dynamics and regulators of cell fate decisions are revealed by pseudotemporal ordering of single cells.” Nature Biotechnology 32 (4): 381–86.

Trevino, Alexandro E., Fabian Müller, Jimena Andersen, Laksshman Sundaram, Arwa Kathiria, Anna Shcherbina, Kyle Farh, et al. 2021. “Chromatin and gene-regulatory dynamics of the developing human cerebral cortex at single-cell resolution.” Cell 184 (19): 5053–69.e23.

Vandereyken, Katy, Alejandro Sifrim, Bernard Thienpont, and Thierry Voet. 2023. “Methods and applications for single-cell and spatial multi-omics.” Nature Reviews. Genetics 24 (8): 494–515.

Wang, Yang, Peng Yuan, Zhiqiang Yan, Ming Yang, Ying Huo, Yanli Nie, Xiaohui Zhu, Jie Qiao, and Liying Yan. 2021. “Single-Cell multiomics sequencing reveals the functional regulatory landscape of early embryos.” Nature Communications 12 (1): 1–14.

Weiler, Philipp, Marius Lange, Michal Klein, Dana Pe’er, and Fabian Theis. 2024. “CellRank 2: Unified fate mapping in multiview single-cell data.” Nature Methods 21 (7): 1196–1205.

Weinreb, Caleb, Samuel Wolock, Betsabeh K. Tusi, Merav Socolovsky, and Allon M. Klein. 2018. “Fundamental limits on dynamic inference from single-cell snapshots.” Proceedings of the National Academy of Sciences of the United States of America 115 (10): E2467–76.

Witoelar, Aree, Arvid Rongve, Ina S. Almdahl, Ingun D. Ulstein, Andreas Engvig, Linda R. White, Geir Selbæk, et al. 2018. “Meta-analysis of Alzheimer’s disease on 9,751 samples from Norway and IGAP study identifies four risk loci.” Scientific Reports 8 (1): 18088.

Wolf, F. Alexander, Philipp Angerer, and Fabian J. Theis. 2018. “SCANPY: Large-scale single-cell gene expression data analysis.” Genome Biology 19 (1): 15.

Wu, Hulin, Tao Lu, Hongqi Xue, and Hua Liang. 2014. “Sparse additive ordinary differential equations for dynamic gene regulatory network modeling.” Journal of the American Statistical Association 109 (506): 700–716.

Xing, Qiao Rui, Chadi A. El Farran, Ying Ying Zeng, Yao Yi, Tushar Warrier, Pradeep Gautam, James J. Collins, et al. 2020. “Parallel bimodal single-cell sequencing of transcriptome and chromatin accessibility.” Genome Research, July. 10.1101/gr.257840.119.

Xu, Wei, Weilong Yang, Yunlong Zhang, Yawen Chen, Ni Hong, Qian Zhang, Xuefei Wang, et al. 2022. “ISSAAC-Seq enables sensitive and flexible multimodal profiling of chromatin accessibility and gene expression in single cells.” Nature Methods 19 (10): 1243–49.

Yan, Rui, Chan Gu, D. You, Zhongying Huang, Jingjing Qian, Qiuyun Yang, Xin Cheng, et al. 2021. “Decoding dynamic epigenetic landscapes in human oocytes using single-cell multi-omics sequencing.” Cell Stem Cell 28 (9): 1641–56.e7.

Zhang, Bing, and Ya-Chieh Hsu. 2017. “Emerging roles of transit-amplifying cells in tissue regeneration and cancer.” Wiley Interdisciplinary Reviews. Developmental Biology 6 (5). 10.1002/wdev.282.

Zhang, Bing, Pai-Chi Tsai, Meryem Gonzalez-Celeiro, Oliver Chung, Benjamin Boumard, Carolina N. Perdigoto, Elena Ezhkova, and Ya-Chieh Hsu. 2016. “Hair follicles’ transit-amplifying cells govern concurrent dermal adipocyte production through Sonic Hedgehog.” Genes & Development 30 (20): 2325–38.

Zhang, Xueying, Rui Gao, Changteng Zhang, Yi Teng, Hai Chen, Qi Li, Changliang Liu, et al. 2023. “Extracellular RNAs-TLR3 signaling contributes to cognitive impairment after chronic neuropathic pain in mice.” Signal Transduction and Targeted Therapy 8 (1): 292.

Zheng, Shijie C., Genevieve Stein-O’Brien, Leandros Boukas, Loyal A. Goff, and Kasper D. Hansen. 2023. “Pumping the brakes on RNA velocity by understanding and interpreting RNA velocity estimates.” Genome Biology 24 (1): 246.

Zhu, Chenxu, Miao Yu, Hui Huang, Ivan Juric, Armen Abnousi, Rong Hu, Jacinta Lucero, M. Margarita Behrens, Ming Hu, and Bing Ren. 2019. “An ultra high-throughput method for single-cell joint analysis of open chromatin and transcriptome.” Nature Structural & Molecular Biology 26 (11): 1063–70.

Zhu, Chenxu, Yanxiao Zhang, Yang Eric Li, Jacinta Lucero, M. Margarita Behrens, and Bing Ren. 2021. “Joint profiling of histone modifications and transcriptome in single cells from mouse brain.” Nature Methods 18 (3): 283–92.

